# Super-CUT&Tag: A Sensitive, Spatially Resolved Approach for Epigenomic Profiling in Tissue Sections

**DOI:** 10.64898/2025.12.08.692886

**Authors:** Ming Zhu, Zhicheng Zha, Jian Xuan, Hanyu Zhao, Jinxian Dai, Jiamao Liu, Youyuan Xia, Hongli Lin, Yiding Yang, Fei Chen, Shichao Lin, Xuzhu Liu, Lingling Wu, Zhi Zhu, Jialiang Huang, Huimin Zhang, Chaoyong Yang

## Abstract

Spatial profiling of protein-DNA interactions is essential for understanding gene regulatory processes, but current methods suffer from low sensitivity, limited resolution, and complex workflows. Here we introduce Super-CUT&Tag (<u>S</u>olid-phase interface embedded with an <u>U</u>ltra-dense DNA barcode array to <u>P</u>rofile <u>E</u>pigenomic landscapes for spatially <u>R</u>esolved CUT&Tag), a robust spatial epigenomic method that directly transfers in situ protein-DNA interactions from tissue sections onto a pre-fabricated spatially barcoded array. Incorporating 3D polyamidoamine (PAMAM) dendrimers markedly improves chromatin capture efficiency, achieving several-fold higher sensitivity compared to current spatial CUT&Tag approaches. Using Super-CUT&Tag, we profile H3K27ac spatial landscape and uncover previously hidden active-chromatin heterogeneity across mouse embryonic tissues and stages. Integration with spatial transcriptomics further reveals spatiotemporal enhancer regulation dynamics of *Neurod2* during corticogenesis. By overcoming key bottlenecks in sensitivity, resolution and usability, Super-CUT&Tag establishes a powerful platform for spatial epigenomic profiling in complex tissues.

## Introduction

Cellular fate determination and gene expression specificity are regulated by diverse epigenetic regulatory mechanisms^1^. Histone modifications are key regulators of spatiotemporal gene expression and play important functions during cell and tissue development^2–4^. These post-translational modifications mediate differential gene activation/silencing through chromatin structural remodeling. For example, H3K27ac (acetylation) is associated with chromatin accessibility and transcriptional activation^5,6^, and H3K4me3 (trimethylation) is associated with activate promoters, whereas H3K27me3 is associated with chromatin condensation and gene repression^7–9^. The balance of these modifications determines the chromatin accessibility landscapes and the transcriptional program, and finally determines the cellular differentiation trajectory^10,11^.

To investigate chromatin components, such as histone modifications and transcription factors, researchers have developed various methodologies such as chromatin immunoprecipitation sequencing (ChIP-seq)^12^, Cleavage Under Targets and Release Using Nuclease (CUT&RUN)^13^, and Cleavage Under Targets and Tagmentation (CUT&Tag)^14^. Among these, CUT&Tag pairs target-specific antibodies with a protein A–Tn5 transposase to cleave and tag DNA *in situ*, offering improved sensitivity and a better signal-to-noise ratio. Its compatibility with single-cell technologies allows researchers detect chromatin components at single cell resolution^15–21^. However, single-cell technologies require cell dissociation leading to the loss of spatial information^22,23^. Consequently, dissociation-based single-cell methods erase the positional context that is essential for understanding how neighboring cells or tissue microenvironments shape chromatin states and gene regulation. Without spatial coordinates, it becomes impossible to pinpoint where epigenetic modifications are remodeled to drive cell fate decisions and orchestrate the spatiotemporal evolution of tissues.

Recent advances in spatial omics have overcome these limitations through innovative technical approaches^24^. Epigenomic MERFISH allows for *in-situ* detection of histone modifications by targeting them with antibodies and inserting T7 promoters using pA-Tn5, followed by converting DNA into RNA and detecting RNA signals with MERFISH^25,26^. Although Epigenomic MERFISH works well for detecting short DNA sequences, it suffers from technical limitations including incompatibility with transcriptomic profiling and technically complexity. Moreover, because Epigenomic MERFISH relies on hybridization-based detection of specific loci, it is unsuitable for genome-wide profiling with limited throughput, and depends heavily on prior knowledge for probe design. Spatial-CUT&Tag platform developed by Deng et al^27^ achieves spatially resolved profiling of histone modifications by first using pA-Tn5 transposase to insert sequencing adapters at antibody-labeled chromatin regions. Spatial barcodes are then delivered directly into tissue via a microfluidic system and ligated to the inserted adapters, thereby encoding spatial information. This elegant strategy enables epigenomic mapping in the tissue section and is compatible with multi-omics extensions^28,29^. However, the complexity and heterogeneity of the tissue structure, such as the presence of cavities, may lead to leakage, blockage, or uneven barcode distribution, thereby reducing the success rate and reproducibility of the experiment. In addition, reliance on microfluidic devices increases technical complexity and hinders large-scale application.

Here, we introduce Super-CUT&Tag, a streamlined and robust spatial epigenomics workflow that combines a solid-phase, ultra-dense DNA-barcode array with 3D PAMAM dendrimers to achieve high-efficiency chromatin DNA capture. Pre-fabricated slides permit quality control before tissue placement, markedly reducing experimental failure and enhancing reproducibility. Compared with Spatial-CUT&Tag, Super-CUT&Tag yields several-fold increase in sensitivity for H3K27ac profiling. We applied this platform to mouse embryos from e10.5 to e14.5 and identified region-specific enhancers that demarcate developmental compartments. At 15 µm resolution, we successfully resolved the heterogeneity of H3K27ac modifications across distinct layers of the mouse forebrain cortex. Integrating spatial epigenomes with matched transcriptomes further uncovered the spatiotemporal circuitry by which Neurod2 governs pallial neuronal identity.

## Results

### Design and overview of Super-CUT&Tag

To achieve highly sensitive spatial profiling of chromatin components, we engineered the Super-CUT&Tag capture slides by attaching fifth-generation (G5) PAMAM dendrimers. Each spherical dendrimer, bearing 128 active amino groups, introduced substantial nanoscale roughness compared with planar (2D) surfaces^30^. This architecture facilitated covalent cross-linking of high-density spatial DNA barcodes and promoted efficient capture of DNA fragments (Fig. 1a). Next, we added unique spatial barcodes onto the aminated surface through a two-step microfluidic process that delivered barcode sets along the x and y axes separately (Supplementary Table 1, 2). The ligation at each intersection produced a dense, combinatorial two-dimensional matrix of spatial barcodes, effectively encoding positional information directly onto the slide surface (Fig. 1a, Supplementary Fig.1). Fluorescence imaging confirmed that G5 PAMAM-modified 3D slides exhibited substantially higher densities of modified DNA barcodes compared with 2D slides (Fig. 1b, c). The spatial DNA barcode density on the 3D dendrimeric slides was calculated to be ∼2.07 × 10^4^ per μm², approximately an order of magnitude higher than that on 2D slides (∼1.16 × 10³ DNA/μm²), resulting in a marked increase in DNA capture efficiency (Fig. 1d, e). Fluorescence imaging further showed strong and uniform barcode signals across 3D dendrimeric DNA arrays, demonstrating highly stable encoding and ensuring robust reproducibility and performance (Supplementary Fig.1d, e). In addition, we replaced the conventional terminal poly(T) sequences on barcodes with sequence-specific spacers, designed to enhance hybridization specificity with tagged DNA fragments via splint oligonucleotides mediation (Supplementary Table 3).

**Fig. 1.**
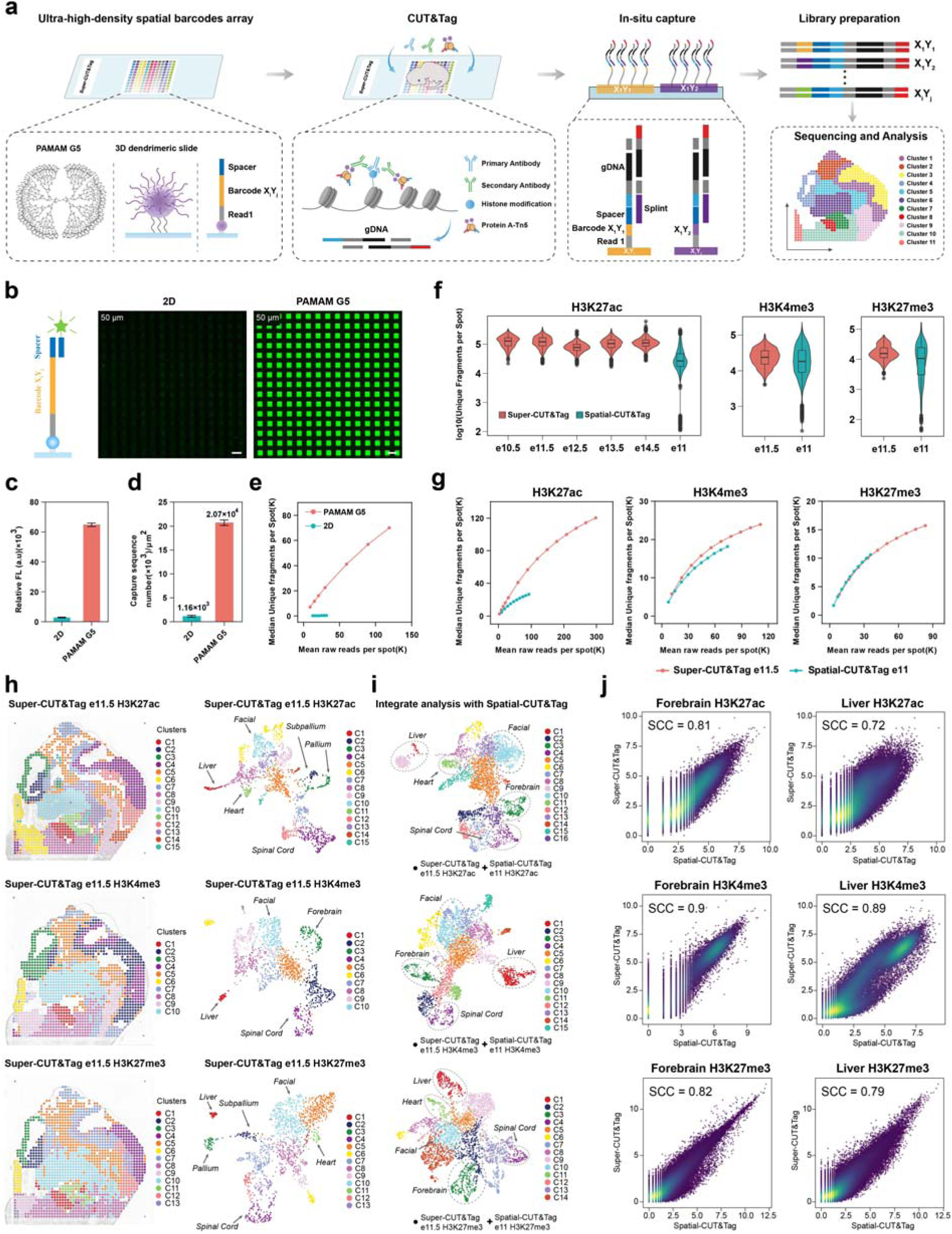
Development and benchmark of Super-CUT&Tag. **a,** Schematic overview of the Super-CUT&Tag workflow for spatial profiling of chromatin modifications. **b,** Fluorescence images of 2D DNA arrays and 3D dendrimeric DNA arrays generated with PAMAM G5; spot size, 50□μm; scale bar, 100□μm, c-d, Comparison of relative fluorescence intensity (**c**) and spatial DNA modification density (**d**) between 2D home-made amino slides and PAMAM G5 dendrimer-coated slides. **e,** Saturation curves showing the median unique fragments per spot versus mean raw reads per spot for Super-CUT&Tag using 2D slides and PAMAM G5 dendrimer-coated slides. f, Comparison of unique fragments per spot across histone modifications (H3K27ac, H3K4me3 and H3K27me3) between Super-CUT&Tag and Spatial-CUT&Tag at a 50□μm resolution. g, Saturation curves comparing Super-CUT&Tag (e11.5) and Spatial-CUT&Tag (e11) across three histone modifications at a 50□μm resolution. h, Spatial distribution (left) and UMAP embeddings (right) of unsupervised clustering based on H3K27ac, H3K4me3 and H3K27me3 profiles in e11.5 embryos. i, Integrated UMAP of chromatin profiles from Super-CUT&Tag (e11.5) and Spatial-CUT&Tag (e11) across three histone modifications. j, Correlation analysis of histone modification signals between Super-CUT&Tag and Spatial-CUT&Tag in forebrain and liver samples across H3K27ac, H3K4me3 and H3K27me3. Spearman correlation coefficients (Spearman’s ρ) are shown to indicate the consistency of peak matrix.

Once the barcoded array is prepared, fresh-frozen (FF) tissue cryosections are directly mounted onto the surface and lightly crosslinked with 0.2% formaldehyde, to preserve both chromatin structures and tissue morphology. CUT&Tag labeling was then performed *in situ* using primary antibodies targeting specific chromatin components, such as histone modifications, followed by secondary antibodies conjugated to a protein A-Tn5 fusion enzyme. A fluorescently labeled secondary antibody was also included to generate images for the morphological reference (Extended Data Fig. 1a). Following Tn5-mediated tagmentation of DNA near the target modification, splint oligo enabled specific hybridization of tagged DNA fragments to spatial barcodes on the slide (Extended Data Fig. 1b, c, Supplementary Table 3). During this step, proteinase K treatment simultaneously reversed crosslinks, allowing DNA to access and interact with the surface-bound DNA barcodes. Also, polyethylene glycol 8000 (PEG8000) was added to the hybridization buffer to limit the diffusion and preserve spatial fidelity (Extended Data Fig. 2a-d). The final spatial barcoding step involved enzymatic ligation and polymerase extension, integrating both spatial barcodes and CUT&Tag-derived DNA fragments into continuous DNA molecules. Subsequent PCR amplification and high-throughput sequencing allowed for decoding of spatial coordinates and DNA fragment sequences, enabling reconstruction of spatially resolved chromatin landscapes across tissue sections.

### Benchmarking of Super-CUT&Tag

To benchmark the data quality of Super-CUT&Tag, we systematically examined its performance in capturing the dynamic landscape of H3K27ac across mouse embryogenesis covering embryonic days 10.5 to 14.5. Additionally, we profiled H3K27me3 and H3K4me3 at e11.5 to assess Super-CUT&Tag’s versatility in capturing distinct histone modifications. Comparative analysis with Spatial-CUT&Tag^27^ at a spatial resolution of 50□μm revealed that Super-CUT&Tag achieved 3-5-fold increase in median unique fragment counts for H3K27ac compared with e11 section detected by Spatial-CUT&Tag (77,274-128,792 vs 26,811; Fig. 1f, g). We merged peaks from H3K27ac ChIP-seq data of different embryonic tissues from the ENCODE^31^ database as unified peaks, 11.9-16.1% of fragments were mapped to these peaks (Supplementary Table 4), slightly outperforming Spatial-CUT&Tag (11.1%) (Extended Data Fig. 2e). For H3K27me3, Super-CUT&Tag yielded a median of 15,656 unique fragments per spot, with 34.6% of reads mapping to ENCODE peaks, comparable to Spatial-CUT&Tag (10,695 fragments per spot and 35.7% in peaks). For H3K4me3, Super-CUT&Tag achieved a higher median fragment count (23,990 vs 18,320) and markedly improved peak enrichment (63.4% vs 35.7%). For all three histone modifications, the fragment length distributions displayed distinct mono- and sub-nucleosomal peaks, confirming the precision and specificity of Super-CUT&Tag in mapping chromatin features (Extended Data Fig. 2f, g). Quality control analyses further validated the Super-CUT&Tag mouse embryo data, confirming consistently high quality and reproducibility across samples (Supplementary Fig. 2).

To evaluate the capability of Super-CUT&Tag in resolving spatial heterogeneity, we performed unsupervised clustering independently on the spatial profiles of H3K27ac, H3K27me3 and H3K4me3. Remarkably, all these histone modifications yielded highly concordant clustering patterns that aligned with anatomical structures across the embryo. For example, clusters C1, C2 and C3 consistently mapped to the liver, subpallium, and pallium regions, respectively. Cluster C4 was enriched in the spinal cord, C10 corresponded to the facial region, and C11 (detected in H3K27ac and H3K27me3 datasets) aligned with the heart (Fig. 1h).

To further dissect inter-clusters, we computed the gene score reflecting the signal enrichment in the gene promotor region. H3K4me3 and H3K27ac gene scores of marker genes were positively correlated, while both showed an inverse relationship with the repressive mark H3K27me3 (Extended data Fig. 3a). For example, the liver marker gene *Gfi1b* and heart marker gene *Hand2* displayed strong H3K4me3 and H3K27ac enrichment, along with low levels of H3K27me3 signals in the liver region and heart region (Extended data Fig. 3b, c). In contrast, the neurogenic marker gene *Ascl1* and *Sox2* exhibited a similarly active chromatin landscape in both the brain and spinal cord (Extended data Fig. 3d, e).

We next integrated Super-CUT&Tag H3K4me3, H3K27me3 and H3K27ac datasets with corresponding Spatial-CUT&Tag data^27^ to validate the accuracy and consistency. After correcting for batch effects by harmony^32^ (Supplementary Fig. 3), we discovered total 16 distinct cell populations. These clusters matched up spatially with specific embryonic tissue regions, with spots from the same anatomical areas clustering together across different histone modification datasets (Fig. 1i). Notably, Super-CUT&Tag exhibited more consistent clustering with fewer scattered points, probably because it had more uniform fragment counts and fewer failures across the tissue (Extended Data Fig. 4a-c). Correlation analysis with both Spatial-CUT&Tag datasets (Fig. 1j) and ENCODE ChIP-seq datasets (Extended Data Fig. 4d) revealed strong concordance, with high spearman correlation coefficients across forebrain and liver of e11.5 embryos. These results established that Super-CUT&Tag offers highly specific and sensitive performance for spatial epigenomic profiling.

To assess the reproducibility of Super-CUT&Tag, we conducted replicate experiments targeting three distinct histone modifications (H3K27ac, H3K4me3 and H3K27me3). UMAP results showed all three histone modifications mapped well even without correcting for batch effects (Supplementary Fig. 4), which indicates that Super-CUT&Tag is quite reliable and produces reproducible results. (Extended Data Fig. 5a). Unsupervised clustering and spatial visualization further confirmed that the same groups of regions were consistently identified and located in the same tissue areas in each replicate (Extended Data Fig. 5b, c), with a strong spearman correlation (spearman’s ρ > 0.92) (Extended Data Fig. 5d). Together, by using pre-fabricated, highly sensitive slides, Super-CUT&Tag enables accurate and reproducible profiling of the spatial epigenome across various histone modifications.

### Spatial H3K27ac Landscapes of Mouse Embryo Development

To explore how active enhancer function during embryonic development, we profiled the spatial distribution of H3K27ac modifications across mouse embryo sections from e10.5 to e14.5 stages and obtained high-quality data (Extended Data Fig. 6a, b). Clustering analysis of each developmental stage revealed 8 to 14 distinct spatial clusters (Fig. 2a). By combining with single-cell RNA sequencing data^33^, cell types matched well with known tissue structures (Fig. 2b, c). For example, at e11.5, radial glial cells were predominantly found in the neural tube, oligodendrocyte precursor cells and inhibitory interneurons were clustered in the subpallium, and developing oligodendrocytes appeared in the pallium. Notably, radial glia and oligodendrocyte progenitors showed a progressive spatial expansion in forebrain and spinal cord regions, while cardiac muscle and erythroid lineages exhibited strong but spatially confined acetylation signals in the heart and hematopoietic niches, respectively (Fig. 2c). These findings underscore the ability of Super-CUT&Tag to capture dynamic epigenomic landscapes in a spatially resolved, cell type–specific manner.

**Fig. 2.**
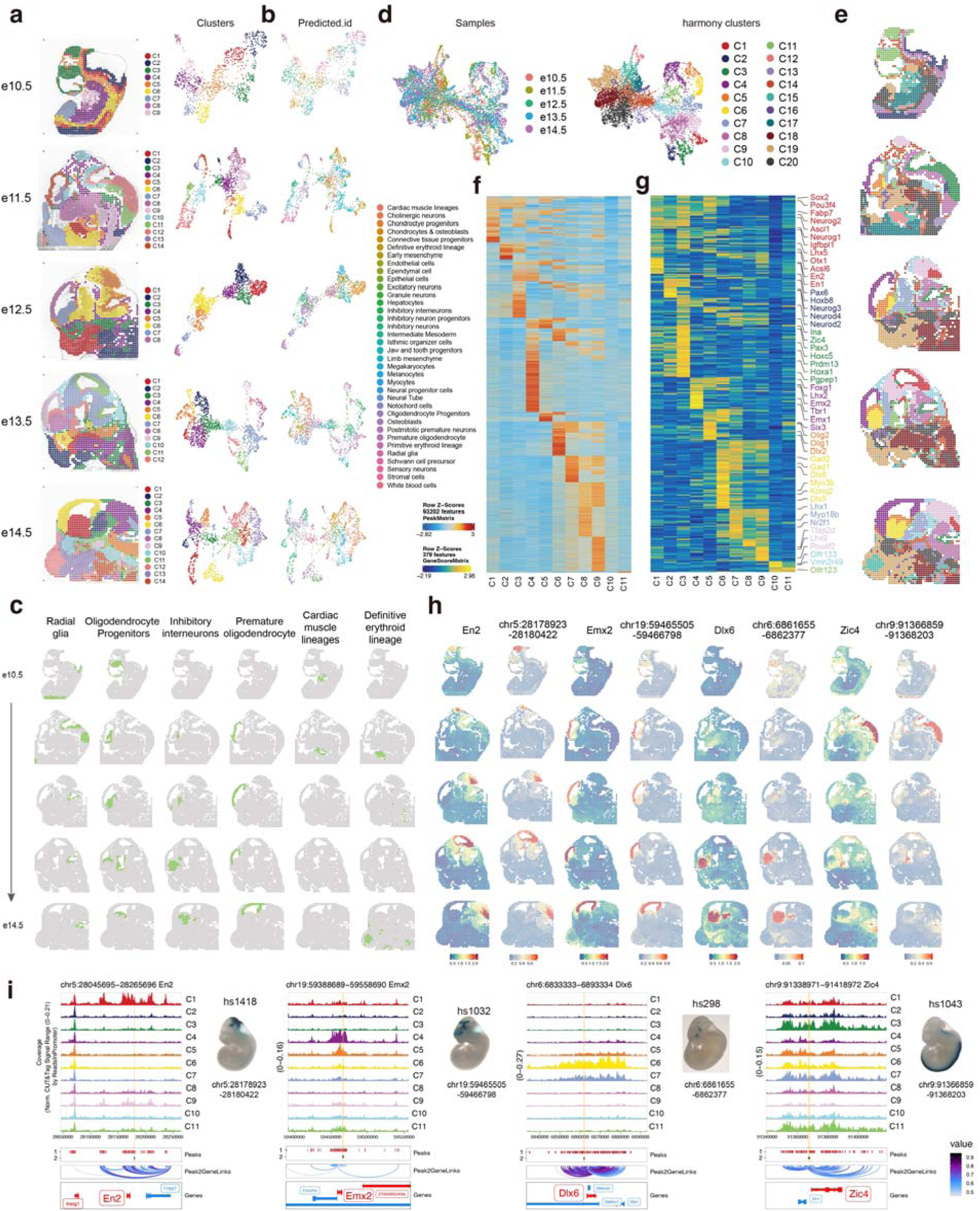
Spatial landscape of H3K27ac modification during mouse embryo development. **a,** Unsupervised clustering of mouse embryo sections for Super-CUT&Tag. **b,** UMAP visualization of cell types annotated with reference scRNA-seq data. **c,** Spatial distribution of cell types annotated with reference. **d,** UMAP visualization of samples (left) and clusters (right) after batch effector removal with harmony. **e,** Spatial visualization of clusters after batch effector removal. **f-g,** Heatmap of marker peaks (**f**) and marker gene scores (**g**) of clusters C1-C11. **h,** Spatial distribution of marker gene associated VISTA enhancers, indicated by gene activity scores (left) and H3K27ac signals (right). **i**, Genome browser plots of H3K27ac signals around. Peaks 1: Super-CUT&Tag peaks; Peaks 2: VISTA enhancer, with their validated enhancer activity measured by VISTA (right).

After correcting for batch effects and integrating chromatin profiles across five developmental stages (e10.5 to e14.5), unsupervised clustering revealed 20 distinct spatial clusters (Fig. 2d, e). Peak calling on the combined dataset yielded 343,966 uniformly sized (501 bp) peaks, excluding those located on mitochondrial or sex chromosomes. Using ArchR^33^, we computed cluster-specific marker peaks and gene scores, enabling high-resolution spatial annotation of embryonic cell populations (Fig. 2f, g, Extended Data Fig. 6c, d). For example, clusters C1 to C11 were predominantly localized to the central nervous system. With forebrain subregions, C4 mapped to the pallium, marked by expression of *Emx2* and *Neurod2*; C5 enriched with *Fabp7*, *Foxn3*, *Hes6* mapped to the subpallium ventricular zone; and C6 mapped to GABAergic neurons (*Dlx1*, *Dlx5*) (Extended Data Fig. 6e). Clusters C9 and C1 captured the midbrain regions, indicated by markers like *Lhx9* and *Tfap2b*, and radial glia markers such as *Dbx1* and *Efna5*. C7 was associated with the diencephalon marked by *Lhx1* activation at e13.5. Clusters C2, C3 and C8 represented hindbrain and spinal cord areas, identified by Hoxb8 and Zic4. Beyond the central nervous system (Extended Data Fig. 6f), C12 was corresponded to liver (*Klf1*, *Slc4a1*, *Hba-a2*), C15 to heart and blood vessels (*Hand2*, *Gata4*, *Nkx2-5*), C18 to muscle (*Atp2a1*, *Trpv1*), C19 to facial mesenchyme (*Prrx2*), and C20 to cartilage (*Matn4*, *Foxd1*). Gene Ontology analysis of top marker genes closely matched these anatomical regions, supporting both the spatial precision of our clustering and the accuracy of cell type annotation (Supplementary Fig. 5).

To identify functional enhancers regulating gene expression, we performed peak-to-gene linkage correlation analysis using ArchR and identified of 50,957 peak-to-gene links (Extended Data Fig. 6g). Several of these predicted enhancers overlap with experimentally validated elements from the VISTA Enhancer Browser^34^. For example, enhancer hs1418 (chr5:28178923-28180422), previously shown by reporter assays to drive transcription in the midbrain, exhibited strong H3K27ac enrichment in the corresponding anatomical region in our dataset (Fig. 2h-i). This enhancer displayed high co-accessibility with *En2*, a key transcription factor for midbrain development, suggesting a regulatory interaction. We also discovered putative enhancers linked to key regional identity genes, including *Emx2* (forebrain pallium), *Dlx6* (subpallium) and *Zic4* (spinal cord), all of which showed prominent H3K27ac enrichment, consistent with their spatially restricted activation patterns during development. Genome-wide association studies have identified numerous noncoding risk variants for neurological diseases, yet their functional interpretation remains challenging^35^. By mapping human orthologues of H3K27ac peaks, we observed clustered enrichment of these risk variants in specific neuronal clusters, with psychiatric and cognitive traits enriched in the brain region (clusters C4 and C6-9), while others like ADHD (attention deficit hyperactivity disorder) were specific to midbrain (C9) (Extended Data Fig. 6h, Method). These results underscore the critical role of cluster-restricted regulatory elements in the genetic architecture of neurological disorders, supported by functional annotations from Super-CUT&Tag profiling.

Together, Super-CUT&Tag revealed spatial heterogeneity of H3K27ac across tissues and cell types during embryonic development and identified putative regulatory enhancers associated with gene expression.

### Spatiotemporal Regulation of *Neurod2* Expression

To explore the enhancer networks involved in pallium development, we integrated H3K27ac Super-CUT&Tag data with sci-RNA-seq^36^ and identified 2,245 domains of regulatory chromatin (DORCs)^37^ with strong peak-to-gene associations (Extended Data Fig. 7a). Among them, 372 genes were linked to pallium-specific regulatory peaks (cluster C4) (Fig. 3a), including key forebrain transcription factors such as *Sox2*, *Pax6* and *Neurod2*. Additional analysis identified *Neurod2*, *Neurod1* and *Emx1* as candidate transcription factors in the pallium, each showing strong H3K27ac signal, gene expression and motif activity in this region (Fig. 3c, Extended Data Fig. 7b-e).

**Fig. 3.**
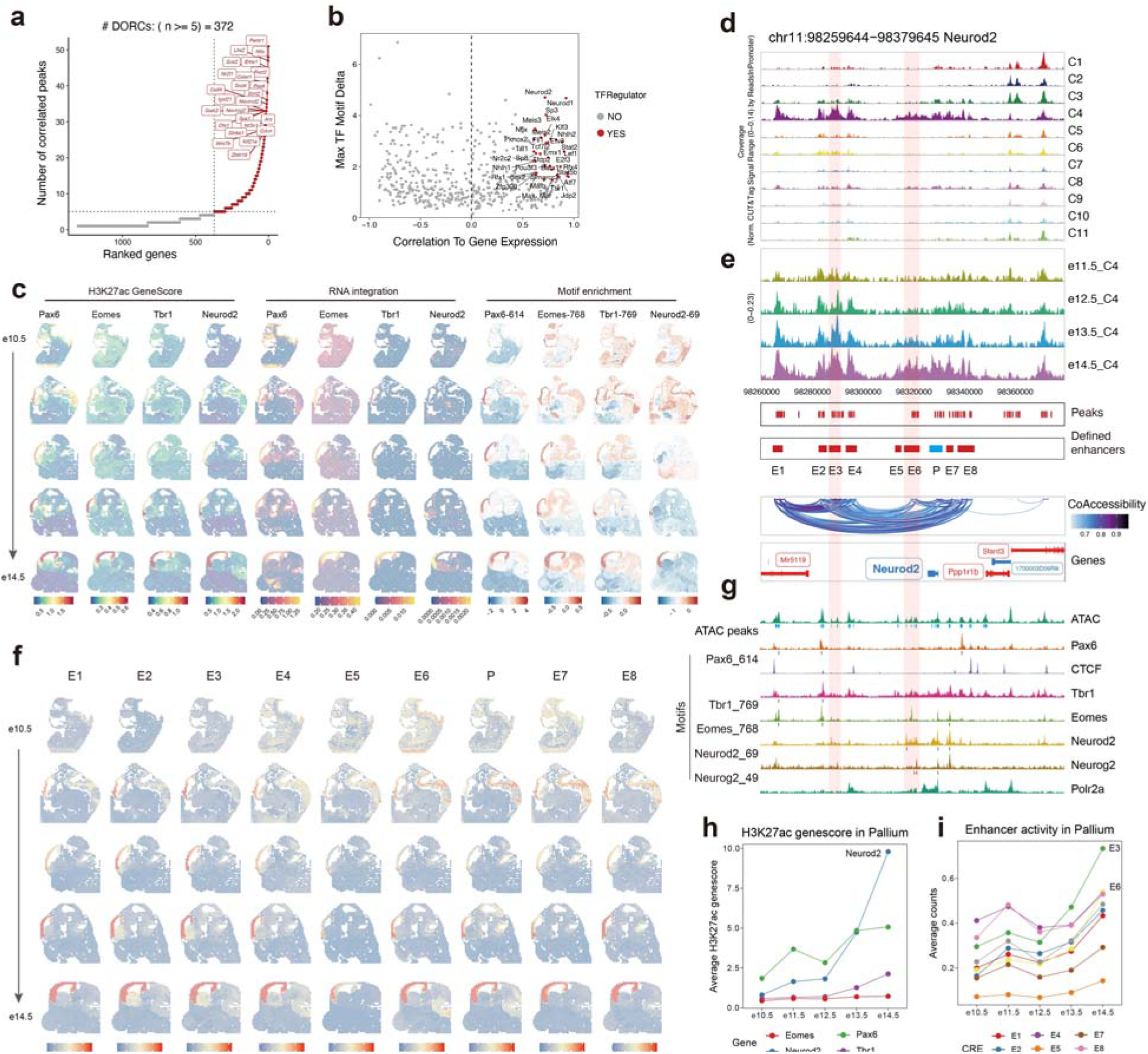
Spatiotemporal regulation of Neurod2 expression by enhancers during embryonic development. **a,** Number of significantly correlated pallium-specific peaks per gene. **b,** Candidate TF regulators in the pallium. **c,** Spatial visualization of H3K27ac gene score (left), integrated RNA expression (middle), and motif deviations (right) for candidate TF regulators. **d-e,** Genome browser tracks of H3K27ac modification across clusters (**d**) and samples (**e**) at the *Neurod2* locus (chr11:98259644-98379645). **f,** Spatial visualization of H3K27ac modification at defined enhancers. **g,** Public ATAC-seq and TF ChIP-seq tracks in neuronal cells at the *Neurod2* locus (chr11:98,259,644-98,379,645). **h,** Average H3K27ac gene score for *Pax6*, *Eomes*, *Tbr1* and *Neurod2* in the pallium across developmental stages. **i**, Average fragment counts (activity) of individual Neurod2-associated enhancers in the pallium across developmental stages.

*Neurod2* (Neurogenic Differentiation Factor 2) belongs to the bHLH (basic helix-loop-helix) family and is a key fate-determining factor in neurogenesis. In the cerebral cortex, *Neurod2* is expressed in excitatory neurons at specific developmental stages, participating in cortical neuronal migration, layering, and morphological maturation^38–41^. A temporally regulated increase in *Neurod2* transcriptional activity was corresponded to a marked gain of H3K27ac signal at its locus from e13.5 onward (Fig. 3c), in line with previous studies^42^. We identified eight candidate enhancers near the *Neurod2* locus by merging adjacent accessible chromatin peaks (Fig. 3d-f), which closely matched ATAC-seq data from e14.5 forebrain in the ENCODE database. Motif enrichment analysis of these enhancers (Extended Data Fig. 7b) suggested potential binding by transcription factors such as *Pax6*, *Eomes* (*Tbr2*) and *Tbr1*, which was further supported by public ChIP-seq datasets in neuron cells (Fig. 3g and Supplementary Table 5). These transcription factors (TFs) are known to function in distinct spatial zones during neocortical development^43^: *Pax6* and *Eomes* are mainly expressed in radial glia within the pallium ventricular zone, while *Tbr1* is enriched in differentiating neurons in the superficial layers. This is consistent with our spatial data, which showed that *Pax6* and *Eomes* were localized to the ventricular zone from e10.5 to e14.5, whereas *Tbr1* was found between the intermediate and marginal zones, with gradually increasing expression. *Tbr1* motif activity was stronger in these regions compared to the ventricular zone, and its H3K27ac gene score profile more closely resembled that of *Neurod2* than *Pax6* (Fig. 3h). Notably, enhancers E3 and E6, bound by *Neurod2* as revealed by ChIP-seq data (Fig. 3g), became highly active after e13.5, suggesting that *Neurod2* may enhance and maintain its own expression by binding to its own regulatory elements (Fig. 3i).

### High-resolution Super-CUT&Tag enables deciphering *Neurod2* spatial enhancer code in the mouse embryonic forebrain cortex

To explore how *Neurod2* expression is spatially regulated, we used a high-resolution spatially barcoded array with 15 μm spots to perform H3K27ac analysis and RNA-seq on neighboring slices from the forebrain pallium of e14.5 mouse embryos. In the Super-CUT&Tag assay, we identified a median of 15,958 fragments across 5,390 spots and 125,170 peaks (uniform 501-bp intervals), with 10.4% of fragments showing overlapping these peak regions (Extended Data Fig. 8a). The clustering analysis of these peaks found 13 clusters (C1-C13, Fig. 4b, Extended Data Fig. 8c). In the spatial RNA-seq assay, we detected a median of 3,622 genes and 7,562 UMIs per spot across 5,554 spots. Clustering analysis identified 20 clusters (R1-R20, Fig. 4c, Extended Data Fig. 8b, c). Both H3K27ac and transcriptome clustering subdivided the forebrain dorsal pallium into three layers: ventricular zone (R4, R16, C5 and C6), intermediate zone (R2 and C11), and superficial layers (R13 and C13). In the ventricular zone, *Eomes* and *Pax6* showed high expression levels, along with higher H3K27ac enrichment. The intermediate zone exhibited highly expression of *Plxna4* and *Cntn2*, while the superficial layer was marked by *Fezf2*, *Slc17a7* and *Tbr1* (Extended Data Fig. 8e, f and Extended Data Fig. 9). Using stereo-seq data^44^ from e16.5 mouse whole brains, we annotated cell types and observed radial glia cells in the ventricular zone, glutamatergic neuroblasts in the intermediate zone, and cortical glutamatergic neurons in the superficial layers (Fig. 4c, Extended Data Fig. 8d).

**Fig. 4.**
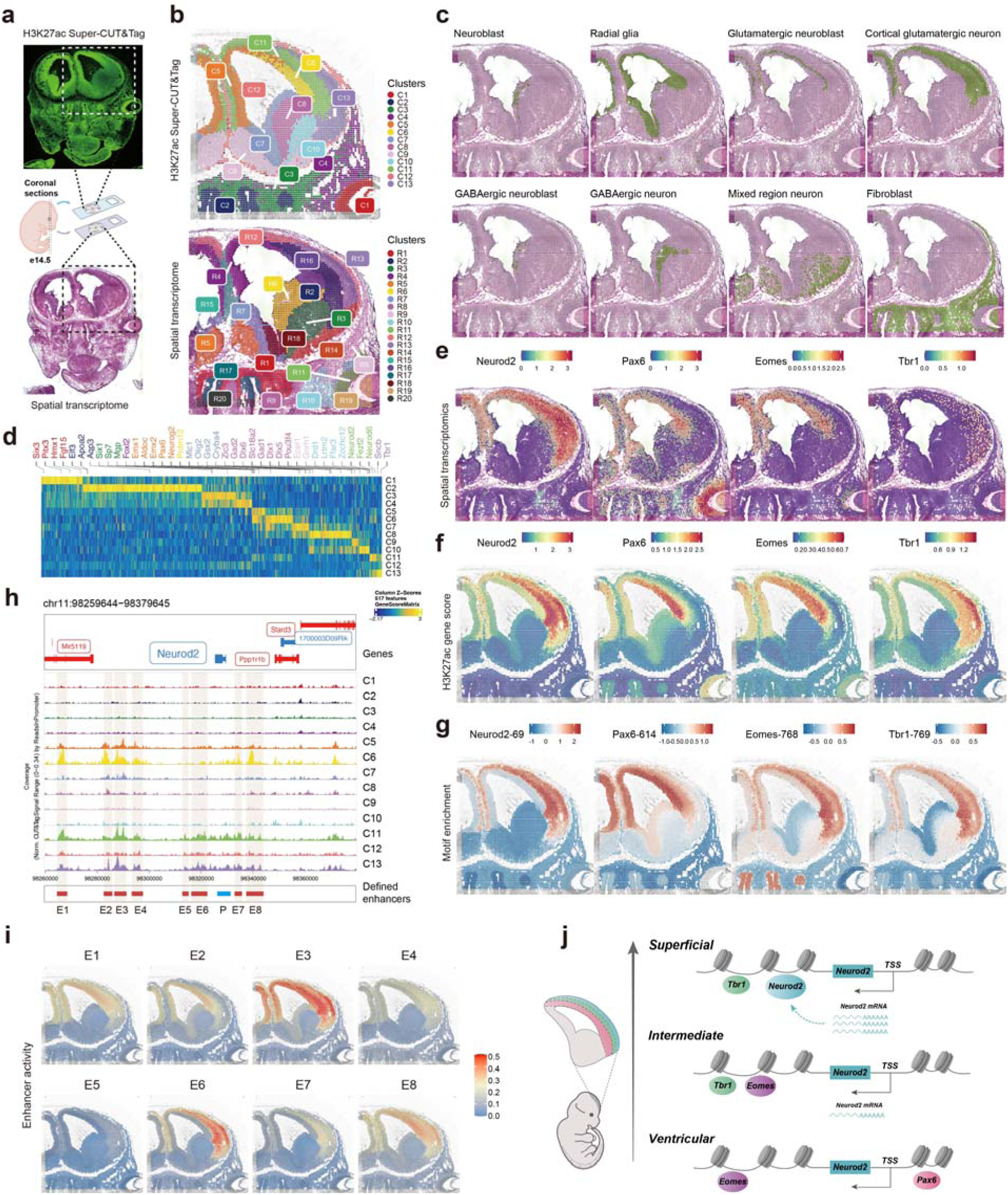
Spatial H3K27ac modification and spatial transcriptome in the e14.5 mouse forebrain. **a**, Schematic of experimental design, in which e14.5 mouse brain tissues were collected and analyzed using H3K27ac Super-CUT&Tag and spatial transcriptomics. **b**, Unsupervised clustering of H3K27ac Super-CUT&Tag data (top) and spatial transcriptomics data (bottom). **c**, spatial distribution of major neural cell types. **d**, Heatmap of marker gene scores of Super-CUT&Tag clusters (C1-C8). **e-g**, Spatial expression patterns (**e**), H3K27ac gene score (**f**) and motif deviations (**g**) of neural development TFs: *Neurod2*, *Pax6*, *Eomes* and *Tbr1*. **h**, Genome browser plots of H3K27ac modification at the *Neurod2* locus across Super-CUT&Tag clusters (C1-C8), showing defined enhancers corelated to *Neurod2*. **i**, Spatial enhancer activity for *Neurod2* enhancers across pallium regions. **j**, Schematic model of *Neurod2* spatial expression regulation.

Next, we explored how the expression of *Neurod2* is regulated by its potential regulatory factors. We observed that *Neurod2* expression levels were quite low in the ventricular zone but gradually increased moving toward the superficial layers (Fig. 4e), where there is strong presence of H3K27ac (Fig. 4f). Motif enrichment analysis indicated potential regulatory functions for *Neurod2* in intermediate zone and superficial layers (Fig. 4g). The spatial distribution of TF regulators of *Neurod2* matches the findings above (Fig. 3c). Specifically, *Pax6* and *Eomes* are abundant in the ventricular zone, while *Tbr1* is expressed in the superficial layers. Further motif analysis revealed that Pax6 mainly acts in the ventricular zone, Eomes functions in both the ventricular and intermediate zones, and Tbr1 is primarily active in the superficial layers (Fig. 4e-g). We also noticed differences in H3K27ac signals at *Neurod2* enhancers: E3 is consistently active across all layers, E1 and E8 show stronger signals in the ventricular zone (C6), E7 (associated with high Eomes ChIP-seq signals) is activated in the intermediate zone, and E6 is highly activated in the intermediate zone and the superficial layers (Fig. 4h, i), where Neurod2 ChIP-seq signal is also detected (Fig. 3g). Then, we selected the spots located in cortex region and calculated the pseudotime to infer the corticogenesis trajectory, and obtained the same pattern using both RNA-seq and H3K27ac data (Extended Data Fig. 10a), which is consistent with the development trajectory of cortical neurons^45^. Along the pseudotime axis, the expression of *Pax6* and *Eomes* gradually decreased from high levels, while the expression of *Tbr1* and *Neurod2* gradually began and continued to increase (Extended Data Fig. 10b, c). Their H3K27ac gene scores also showed the same pattern (Extended Data Fig. 10d, e). The activities of *Neurod2* enhancers exhibited distinct patterns (Extended Data Fig. 10f), with E1 and E2 decreasing along the pseodutime axis, while E5 and E6 showed an increasing trend. E3 remained at a consistently high level, in agreement with earlier observations that it showed the greatest activity at e14.5 (Fig. 3i). Collectively, these findings suggest that *Neurod2* expression is regulated in a flexible way by its proximal enhancers. In the ventricular zone, Pax6 and Eomes act as early regulators to prime *Neurod2* expression. As neurons differentiate, Tbr1 begins to regulate *Neurod2* expression in the intermediate zone. Finally, in the superficial layers, Neurod2 binds to its own enhancer E6, forming an autoregulatory loop that boosts and maintains its activity (Fig. 4j).

By combining Super-CUT&Tag with high-resolution spatially barcoded array, we successfully dissected H3K27ac heterogeneity in complex structures of the forebrain, spatially revealed the differentiation and migration trajectory of cortical neurons, uncovered the spatial regulatory mechanisms governing *Neurod2* expression, and provided new insight into epigenetic regulation during brain development.

## Discussion

Spatially resolved profiling of the epigenome is essential for understanding how chromatin states orchestrate cellular identity and tissue organization during development. While spatial transcriptomics has benefited from user friendly spatially barcoded array-based technologies^30,46,47^ that offer simplified workflows and variable spatial resolutions, similar approaches for chromatin profiling have faced technical hurdles^48,49^. Transcriptome libraries are broadly distributed within the cytoplasm, while chromatin DNA is kept within the nucleus, resulting in limited contact with array surfaces. We hypothesized that the limitation of the contact area with the array leads to saturation of the capture efficiency. As a result, DNA barcoded probe density must be increased for efficient chromatin DNA retrieval. To address these challenges, we developed Super-CUT&Tag, a high-sensitivity spatial epigenomics platform that overcomes key limitations of previous approaches and enables robust profiling of histone modifications with high spatial resolution. By integrating ultra-high-density barcode arrays with 3D PAMAM dendrimer-enhanced surfaces, Super-CUT&Tag achieves significantly improved DNA capture efficiency and signal sensitivity, enabling 3-5 fold increases in H3K27ac detection compared to existing methods. The resulting data detected by Super-CUT&Tag were highly concordant with ENCODE ChIP-seq references and previously published Spatial-CUT&Tag^27^ datasets, validating the accuracy, sensitivity and reproducibility of our approach. Importantly, Super-CUT&Tag maintained high spatial fidelity and reproducibility across replicates without the need for computational batch correction, suggesting strong robustness of the pre-fabricated capture slides and the simplified workflows.

Taking advantage of the efficient detection capability of Super-CUT&Tag for H3K27ac, we generated a spatial atlas of H3K27ac during mouse embryonic development from e10.5 to e14.5. By integrating spatial epigenomic and transcriptomic data, we revealed distinct spatial clusters corresponding to embryonic tissues and organ systems, and identified cell type-specific enhancers. Many predicted enhancers overlapped with experimentally validated VISTA elements, further supporting the functional relevance of our results. Focusing on forebrain development, we dissected the spatial regulation of *Neurod2*, a key neuronal transcription factor. By integrating H3K27ac data with high-resolution RNA-seq, we precisely visualized the migration trajectories of radial glial cells during cortical development and identified a series of important transcriptional regulators, including *Pax6*, *Eomes* and *Tbr1*, that control *Neurod2* expression in a spatially and temporally coordinated manner. Furthermore, we showed that *Neurod2* may maintain its own expression via binding to its distal enhancers, forming a positive feedback loop during corticogenesis. These findings illustrate the power of Super-CUT&Tag to decode spatiotemporal gene regulation and reconstruct enhancer logic in complex developmental contexts.

Compared to existing spatial epigenomic platforms such as Spatial-CUT&Tag^27^ and Epigenomic MERFISH^25^, Super-CUT&Tag offers several advantages. First, it enables genome-wide detection with higher throughput than probe-based imaging strategy. Second, it separates slide preparation and quality control processes, eliminating the need for on-tissue microfluidic devices for barcoding process, which enhances experimental success rates, stability and reproducibility. This user-friendly method also makes it more suitable for widespread adoption, similar to the approach of 10x Genomics. Super-CUT&Tag improves detection sensitivity by increasing barcode diversity through tree-like macromolecular structures, with further enhancements achievable using higher-generation PAMAM. Moreover, Super-CUT&Tag holds strong potential for adaptation to a wider array of epigenetic marks and chromatin-associated proteins like transcription factors.

Despite these strengths, several limitations remain. First, Super-CUT&Tag currently requires cryosectioned tissues, which may limit its application to archival or FFPE (Formalin Fixed Paraffin Embedded) samples without optimization. This limitation may be overcome by optimizing crosslink reversal strategies, refining tissue permeabilization conditions, and drawing on recent advances in CUT&Tag compatible FFPE workflows^50–52^. Second, although we achieved 15-μm resolution, pushing the resolution to the single-cell or subcellular level will require further miniaturization of spatially barcoded arrays, or employment of high-throughput sequencing flow cells for spatial information capture to achieve near-subcellular resolution^44,53^. Third, while our focus here was on histone modifications, extending Super-CUT&Tag to transcription factors or chromatin remodeling complexes may pose additional challenges related to antibody specificity and signal abundance. These challenges can be addressed by developing high-affinity recombinant antibodies or nanobodies to enhance specificity, and optimization of the tag cleavage chemistry to improve sensitivity^17,21,29,54,55^.

Looking forward, Super-CUT&Tag provides a versatile platform for spatial epigenomics with broad applicability in developmental biology, cancer research, and disease modeling. Integration with single-cell transcriptomics, lineage tracing, or spatial proteomics could further enhance our understanding of gene regulatory networks in situ. Moreover, the spatially barcoded array-based strategy used in Super-CUT&Tag could be extended to other spatial omics modalities^28,29,49^, paving the way for truly integrated multi-omics spatial atlases across tissues and organisms.

## Methods

### Animal

All animal experiments conducted in this study complied with ethical regulations for animal research and were approved by the Institutional Animal Care and Use Committee of Xiamen University (Ethical Approval Number: XMULAC20220298). Pregnant C57BL/6J female mice were euthanized via cervical dislocation, and e10.5, e11.5, e12.5, e13.5, and e14.5 embryos were dissected and collected. Embryos were rapidly rinsed with pre-chilled 1× PBS, blotted dry using dust-free paper, and immediately embedded in pre-cooled Tissue-Tek OCT (Sakura, 4583), followed by rapid freezing on dry ice. The samples were subsequently stored at -80°C until further use. Sagittal sections of embryonic tissues were cut into 10-μm-thick slices using a Leica CM1950 cryostat and mounted on slides for Super-CUT&Tag.

### Preparation of 3D dendrimeric slides

The production and preparation of 3D dendrimeric slides followed the published protocol^30^. Glass slides were cleaned and activated with Piranha solution for 2 hours, followed by treatment with a 5% (3-Glycidyloxypropyl) trimethoxysilane (GOPTS) solution in anhydrous ethanol for 5 hours. The slides were then washed with anhydrous ethanol, dried under nitrogen, and baked at 110°C for 1 hour. A 0.3% (v/v) G5 PAMAM dendrimer solution in methanol was prepared, and the silanized slides were immersed in the solution overnight at 55 rpm. After modification, the slides were washed with methanol, dried, and stored at 4°C until use.

### Microfluidic device design and fabrication

Briefly, the computer-aided design (CAD) files for the required microfluidic chip layout were generated and subsequently used to fabricate a chromium photomask (Supplementary Fig.1b). SU-8 photoresist (SU-8 3050, Microchem) was spin-coated onto a silicon wafer, followed by photolithography using the photomask. Polydimethylsiloxane (PDMS) was prepared by mixing base and curing agent at a ratio of 10:1. After pumping out all the air bubbles, PDMS was then poured onto the silicon mold and cured at 125°C for 25min. After curing, the PDMS microfluidic chip was cut out the fabrication.

### Spatially resolved barcoding on slide surfaces

The fabrication of spatially encoded chips for immobilizing spatial DNA barcodes on 3D dendrimeric slides using a two-step reaction. Briefly, barcode x was first immobilized via DSS-mediated conjugation, followed by succinic anhydride blocking to quench residual amino groups. Barcode y was annealed with a complementary ligation linker and subsequently ligated onto the slide using T4 ligase. The DNA barcoding region was then incubated with 80 mM KOH for 30 seconds to cleave the linker, followed by rinsing, drying, sealing, and storage at 4°C.

### Quantification of spatial DNA barcode density on 3D dendrimeric slides

Fluorescence signals were measured using a microplate reader. Three experimental and three control groups were prepared, each containing barcode y labeled with 488 fluorophores. The experimental groups were used to quantify non-specific adsorption of barcode y onto the slide surface, whereas the control groups were used to calculate the number of spatial DNA barcodes covalently immobilized. In the control groups, barcode y was only hybridized with immobilized barcode x in the absence of T4 ligase. To perform fluorescence-based quantification, a standard fluorescence curve was first generated using a concentration gradient (0.25, 0.5, 1.0, 1.5, and 2.5 μM) of barcode y labeled with fluorophore 488. The reaction wells (7 × 7□mm²) on the slides were washed three times with 2× SSC buffer, followed by the addition of 70 μL of 80 mM KOH three times, each incubated for 10 minutes to fully release barcode y hybridized to barcode x. The eluates were then neutralized with 30 μL of Tris-HCl (pH 7.0), and 150 μL of the resulting solution was collected for fluorescence measurements. The actual number of immobilized barcode y molecules was calculated by subtracting the fluorescence signal of the experimental group from that of the control group, and the surface density was subsequently determined using the standard curve.

### DNA barcode sequences, DNA oligos and other key reagents

All DNA barcode sequences are provided in Supplementary Table 1 and 2. DNA oligos used for PCR and library construction are shown in Supplementary Table 3. All other key reagents are listed in Supplementary Table 6.

### Antibodies

Primary antibodies used were H3K4me3 (Active Motif cat. no. 39159, lot: 24117228), H3K27ac (Abcam cat. no. ab177178, lot: 1104872), H3K27me3 (Active Motif cat. no. 39155, lot: 24354142) and Secondary antibody used were Goat Anti-Rabbit IgG H&L (Abcam cat. no. ab6702, lot: 1036471), Goat Anti-Rabbit IgG H&L (Alexa Fluor® 488) (Abcam cat. no. ab150077, lot: GR3429073).

### Super-CUT&Tag workflow

The frozen tissue slide was thawed for 1□min at 37□°C to adhere the tissue to the slide. Then, the tissue section was fixed with 0.2% formaldehyde for 10□min and quenched with 1.25□M glycine for a further 5□min. After fixation, the tissue was blocked with 2% BSA in Dulbecco’s PBS (DPBS) for 10 minutes to prevent nonspecific tissue adsorption. After blocking, tissue was washed twice with wash buffer (20□mM HEPES pH□7.5, 150□mM NaCl, 1× protease inhibitor cocktail, 0.5□mM Spermidine). The tissue section was then permeabilized for 10□min with CA630-digitonin wash buffer (0.02% digitonin, 0.1% CA630 in wash buffer). The primary antibody was diluted 1:50 in antibody buffer (0.02% BSA, 2 mM EDTA in CA630-digitonin wash buffer), then added and incubated at 4°C overnight. The primary antibody was then removed, and secondary antibody (Goat Anti-Rabbit IgG, 1:50 dilution with antibody buffer) was added with incubation for 30□min at room temperature. Unbound antibodies were removed by washing three times with Wash buffer, each for 1 minute. A 1:30 dilution of pA-Tn5 adaptor complex in 300-wash buffer (20□mM HEPES pH□7.5, 300□mM NaCl, 1× protease inhibitor cocktail, 0.5□mM Spermidine) was added with incubation at room temperature for 1□h. Excess pA-Tn5 protein was removed using 300-wash buffer. Tagmentation buffer (10□mM MgCl_2_ in 300-wash buffer) was added with incubation at 37□°C for 1□h. Next, washing was performed three times with 0.1×SSC for 1□min each and the slides were then spin dried and imaged in a Nikon Eclipse Ti2 inverted fluorescence microscope (×10 magnification) to record fluorescent images of the tissue and area fiducials.

To hybridize the DNA fragments with the barcoded surface oligonucleotides, the sections were then incubated with splint oligonucleotide (in 20 mM Tris-HCl pH 8.0 buffer containing 0.1% CA-630, 300 mM NaCl, 5 mM EDTA, 2□mg/ml Proteinase K, and 10% PEG8000) overnight at 37□°C. Next, the sections were washed twice with 2× NEBuffer 2, and subsequently incubated with ligation and polymerization solution (1× NEBuffer 2 containing 15 U of T4 DNA polymerase, 2,000□U of T4 DNA ligase, 500□µM dNTPs, 1□mM ATP) and incubated at 18□°C for 4□h. Tissue removal was carried out using 2.5 mg/ml Proteinase K in buffer PKD followed by incubation at 56°C for 40□min. The sections were then sequentially washed with 2× SSC 0.1% SDS, 0.2× SSC and 0.1× SSC.

Spatially barcoded single-stranded DNA fragments were released from the arrays by denaturation with 0.08□M KOH for 10□min at room temperature, followed by neutralization with 1□M Tris-HCl pH 7.0. The eluted DNA from each sample was amplified with KAPA HiFi Hotstart ReadyMix. The following PCR program was used: 98□°C for 3□min; 6 cycles of 98□°C for 20□s, 63□°C for 20□s and 72□°C for 1□min; The pre-amplified PCR products were purified using 0.9× VAHTS DNA Clean Beads using the standard protocol and eluted in 21□µl of nuclease-free water. To determine additional cycles, ten microliters of quantitative PCR (qPCR) reaction was prepared, consisting of pre-amplified purified product, primers, KAPA qPCR ReadyMix, and nuclease-free water. The reaction was immediately loaded into a qPCR instrument (LightCycler480 II, Roche Diagnostics) and run under the following thermal cycling conditions: 98°C for 3 min, followed by 25 cycles of 98°C for 5 s and 63°C for 30 s. Finally, the pre-amplified product was mixed with the index primers and KAPA HiFi Hotstart ReadyMix), then PCR reaction performed with the following program: 98□°C for 3□min; n cycles of 98□°C for 20□s, 63□°C for 20□s and 72□°C for 1□min. The value of n was determined based on the quantification cycle (C_q_) value obtained from qPCR. The indexed libraries were cleaned up using 0.8× VAHTS DNA Clean Beads and final concentrations were quantified with a Qubit dsDNA assay kit. Next-generation sequencing (NGS) was performed using Illumina NovaSeq 6000 with PE150 mode.

### Spatial transcriptome

FF tissue sections from the mouse forebrain were cut and mounted onto adhesive slides, followed by H&E staining. Library preparation was performed using the DynaSpatial FF Spatial Gene Expression Kit (Dynamic Biosystems, Suzhou, China) according to the User Guide. After quality control, the library is sequenced on the DNBSEQ-T7 platform.

### Data process

Fluorescence-labeled microscopy images of secondary antibodies in tissue cell nuclei were processed using the DynamicST Assist software (https://github.com/DynamicBiosystems/DynamicST-Assist) to identify regions of interest within the tissue. A custom script was then employed to generate a spatial folder containing data analogous to 10x image data, which can be directly loaded using the Read10X_Image function in Seurat^56^ (v5.0.0). The FASTQ files were first processed with a custom script to extract spatial barcodes, followed by analysis with Cell Ranger ATAC (v2.1.0) using a provided whitelist to generate fragment files. Sequencing quality and mapping statistics for all samples are summarized in Supplementary Table 7. These fragment files were subsequently used as input for ArchR^33^ to perform downstream analyses.

Fragment files were processed in ArchR using the createArrowFiles function to generate tile matrices (5 kb resolution) and gene score matrices, followed by ArchRProject construction. Iterative Latent Semantic Indexing (LSI) was performed for data normalization and dimensionality reduction. Batch effects were corrected using Harmony, followed by dimensionality reduction via Uniform Manifold Approximation and Projection (UMAP) using the addUMAP function and initial clustering with addClusters. For comparative analysis, we generated a peak matrix from ENCODE-defined peaks in ArchR as a benchmark against spatial-CUT&Tag data. In the H3K27ac integration analysis, peak calling was performed on initial clusters to generate the peak matrix, which was subsequently used to compute UMAP projections and cluster assignments following the steps above.

### GO enrichment analysis

Markerpeaks of each clusters were found by getMarkerFeatures using PeakMatrix, top markerpeaks were performed enrichment analysis by rGREAT^57^ with “BP” gene set. While markergenescores were found by getMarkerFeatures using GeneScoreMatrix, top markergenescores were performed enrichment analysis by clusterProfiler^58^ with “BP” gene set.

### Motif enrichment

In the motif enrichment analysis, motif annotations were added using the addMotifAnnotations function with motifSet = “cisbp” to identify known transcription factor binding sites. Background peaks were then selected using addBgdPeaks. Motif deviations were calculated using addDeviationsMatrix, which quantifies accessibility variation of each motif relative to the background. For ATAC-seq peaks, the PeakMatrix was regenerated with identified peaks and the motif enrichment process was repeated.

### GWAS enrichment

To enable comparison to GWASs of human phenotypes, we used liftOver with settings ‘-minMatch=0.5’ to convert accessible elements from mm10 to hg19 genomic coordinates^59^. Next, we reciprocal lifted the elements back to mm10 and only kept the regions that mapped to original loci. We further removed converted regions with lengths greater than 1 kb. We obtained GWAS summary statistics for quantitative traits related to neurological disease and control traits. We prepared summary statistics to the standard format for linkage disequilibrium score regression. We used homologous sequences for each major cell types as a binary annotation, and the superset of all candidate regulatory peaks as the background control. For each trait, we used cell-type-specific linkage disequilibrium score regression (https://github.com/bulik/ldsc) to estimate the enrichment coefficient of each annotation jointly with the background control^60^.

### Cis-regulatory associations

We performed cis-regulatory correlation analysis using FigR^61^, with the PeakMatrix and GeneIntegrationMatrix (generated by ArchR) as input datasets for correlation calculations. Genes exhibiting significant correlations with more than 5 peaks were defined as DORCs (domains of regulatory chromatin).

### Data visualization

The ArchR projects were converted into Seurat objects using customized functions adapted from ArchRToSignac^62^ for spatial image plotting.

## Data availability

The raw sequence data reported in this paper have been deposited in the Genome Sequence Archive^63^ in National Genomics Data Center^64^, China National Center for Bioinformation / Beijing Institute of Genomics, Chinese Academy of Sciences (GSA: CRA025994) that are publicly accessible at https://ngdc.cncb.ac.cn/gsa. The processed data and images in this paper have been deposited in the OMIX (https://ngdc.cncb.ac.cn/omix, accession no. OMIX010273). Published data used in this study include mouse organogenesis cell atlas (MOCA)^36^: Spatial-CUT&Tag data: GSM5028434, GSM5028435, GSM5028436. ChIP-seq data and ENCODE data see Supplementary Table 5.

## Code availability

All codes used for data analysis in this study are publicly available on GitHub at https://github.com/zhuming0301/Super-CUT-Tag.

## Acknowledgements

We thank the National Key R&D Program of China (2024YFF1206600), the National Natural Science Foundation of China Grants (22274066, 92474104 and 32370586), and the Natural Science Foundation of Fujian Province of China (2024J010047) and the Fundamental Research Funds for the Central Universities (20720230068) for their financial support.

## Author contributions

M.Z., H.Z., and C.Y. conceived and designed the experiments. M.Z., Z.Z. performed the experiments. M.Z. performed the bioinformatics analysis. H.Z fabricated microfluidic devices. M.Z. and Z.Z. collected the samples. M.Z. and Z.Z. wrote the paper with the input from other authors. All authors read and approved the final version of the manuscript. C.Y., H.Z. and J.H. and revised the paper and supervised the project.

## Competing interests

C.Y. is a co-founder of Dynamic Biosystems. All other authors declare no competing interests.

---

**Extended data Fig. 1.**
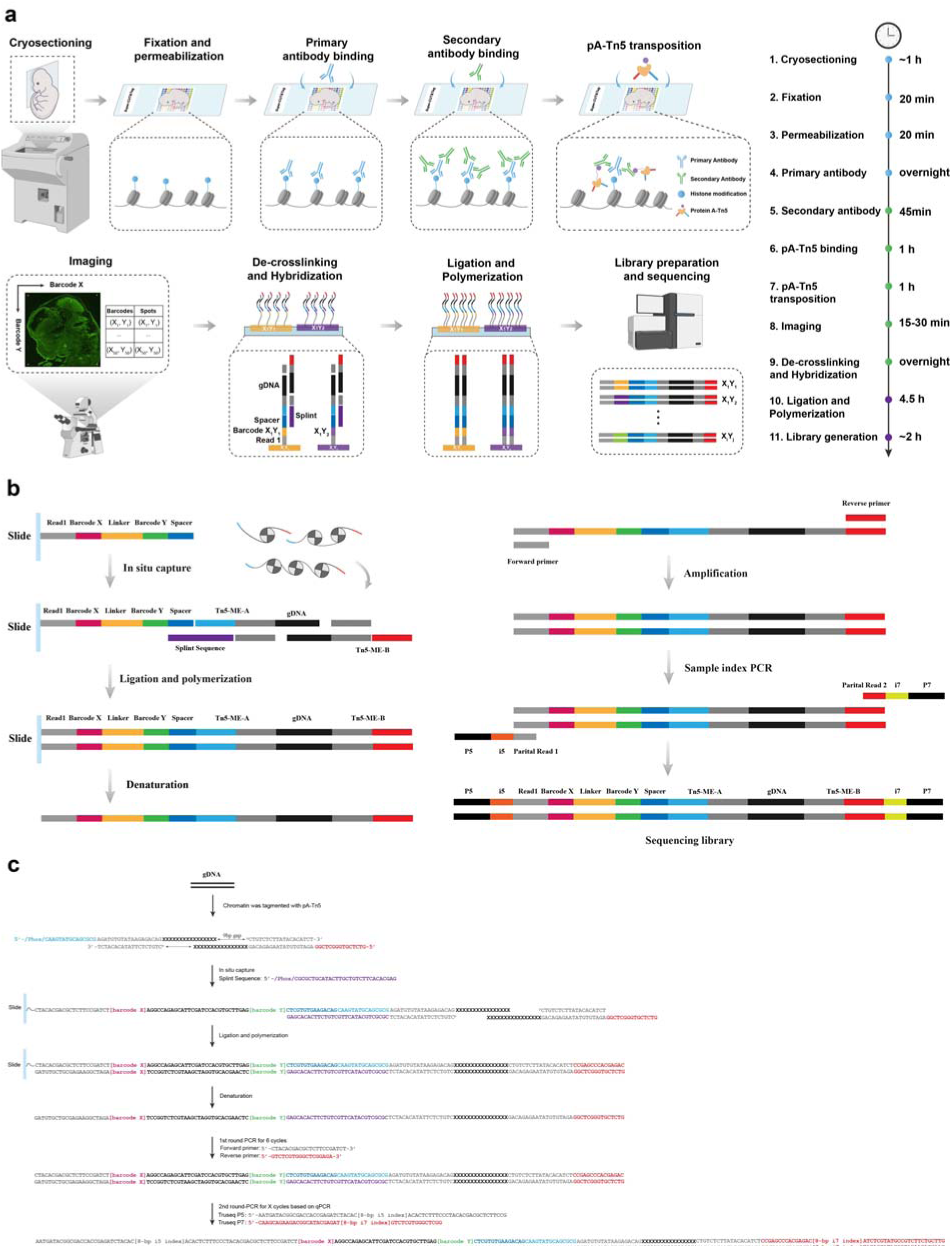
Chemistry workflow and molecular design of Super-CUT&Tag. **a,** Tissue sections were mounted onto Super-CUT&Tag slides, followed by mild formaldehyde fixation and permeabilization to preserve tissue morphology while allowing molecular accessibility. Subsequently, the tissue was incubated with a primary antibody specific to histone modifications or chromatin-associated proteins, followed by the addition of a secondary antibody to enhance anchoring of the pA-Tn5 transposome. Activation of the pA-Tn5 complex was initiated by the addition of Mg²⁺, followed by incubation at 37□°C, thereby facilitating the insertion of predefined adapter sequences at antibody recognition sites on genomic DNA. After imaging to record the spatial coordinates of the tissue, proteinase K was used during library capture to reverse crosslinking, and a splint oligonucleotide was introduced to immobilize DNA fragments onto the slide. Spatial barcoding was then completed through enzymatic ligation and polymerization reactions mediated by ligase and polymerase, generating sequencing-ready libraries. DNA fragments were subsequently eluted using potassium hydroxide, followed by PCR amplification to complete library construction. **b,** A simplified overview of the nucleotide sequences involved in the Super-CUT&Tag library preparation workflow. This schematic highlights the main components and key steps without detailed annotation, serving as a concise counterpart to the more comprehensive illustration presented in c. **c,** Detailed nucleotide sequences involved in each step of the Super-CUT&Tag library preparation workflow are illustrated. Depicted elements include spatial barcode sequences immobilized on the chip, adapter sequences inserted by pA-Tn5 at genomic sites, splint oligonucleotide-mediated hybridization of DNA fragments to the chip surface, and the enzymatic ligation and polymerization steps that link and extend these fragments. The final structure of the sequencing-ready library is also shown, including the arrangement of barcodes and adapter sequences for PCR amplification. Junctions, primer-binding sites, and barcode positions are annotated to illustrate how spatial information is preserved during library construction.

**Extended data Fig. 2.**
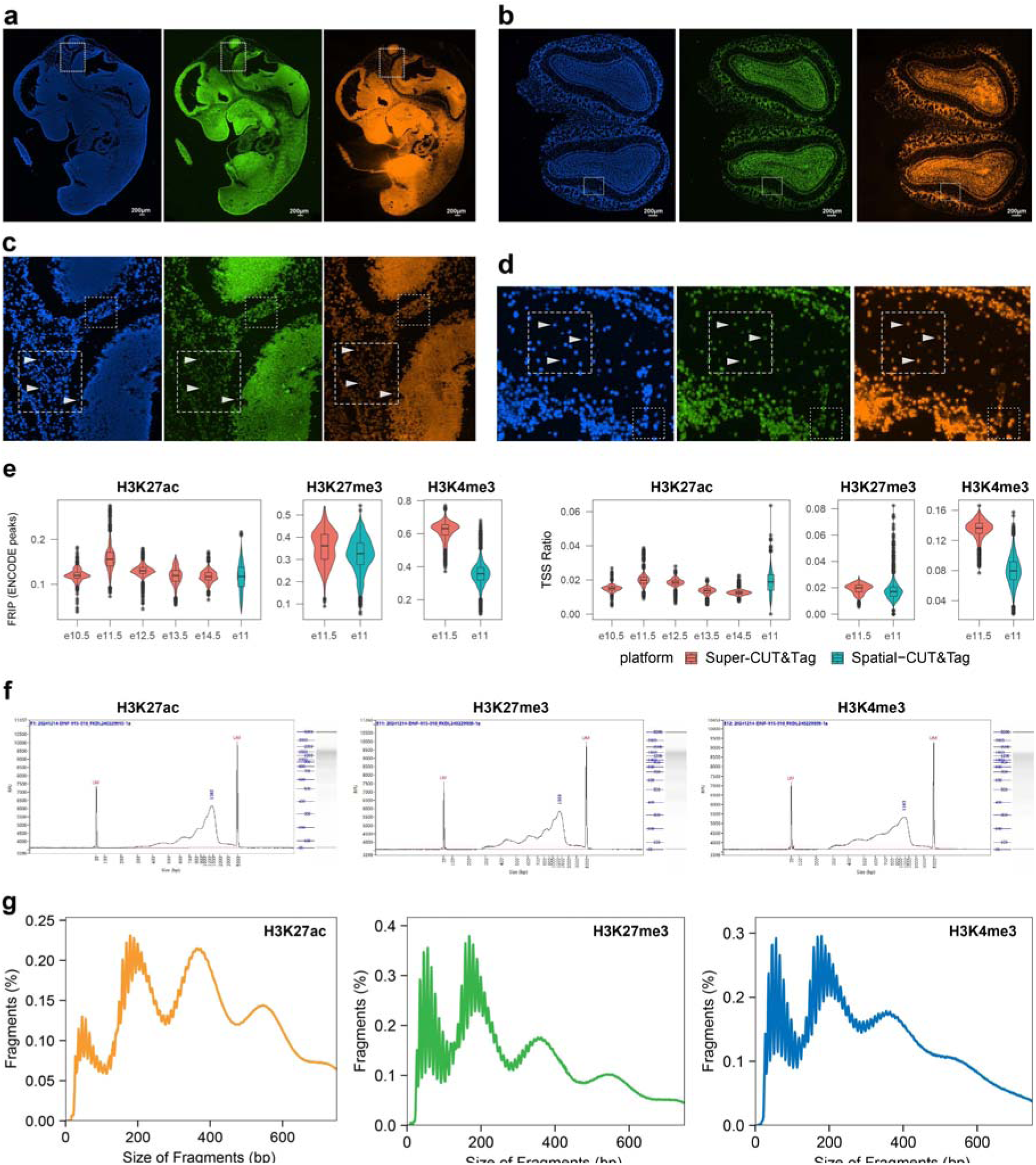
Spatial localization and Quality metrics of CUT&Tag libraries in tissue sections. **a-b,** Fluorescence images of mouse embryonic tissue **(a)** and olfactory bulb tissue **(b)** undergoing the Super-CUT&Tag workflow. Left: DAPI staining of nuclei. Middle: Alexa Fluor 488 signal from the secondary antibody following primary antibody binding to histone modifications or chromatin-associated proteins. Right: Cy3 fluorescence from double-stranded DNA synthesized using Cy3-labeled dCTP after CUT&Tag library capture, tissue removal, and alkaline denaturation. **c-d,** Magnified views of the regions outlined in white dashed boxes in **(a)** and **(b)**, respectively. White triangle arrowheads indicate nuclear positions. The distribution of fluorescent CUT&Tag DNA fragments closely resembled the patterns observed in the DAPI-stained nuclei and Alexa Fluor 488-labeled secondary antibody signals, indicating that the CUT&Tag libraries were spatially confined beneath the cells with minimal diffusion. **e,** Comparison of FRiP scores and TSS enrichment ratios between Super-CUT&Tag and Spatial-CUT&Tag. **f,** Bioanalyzer data of DNA fragments. **g,** Distribution of fragment lengths.

**Extended data Fig. 3.**
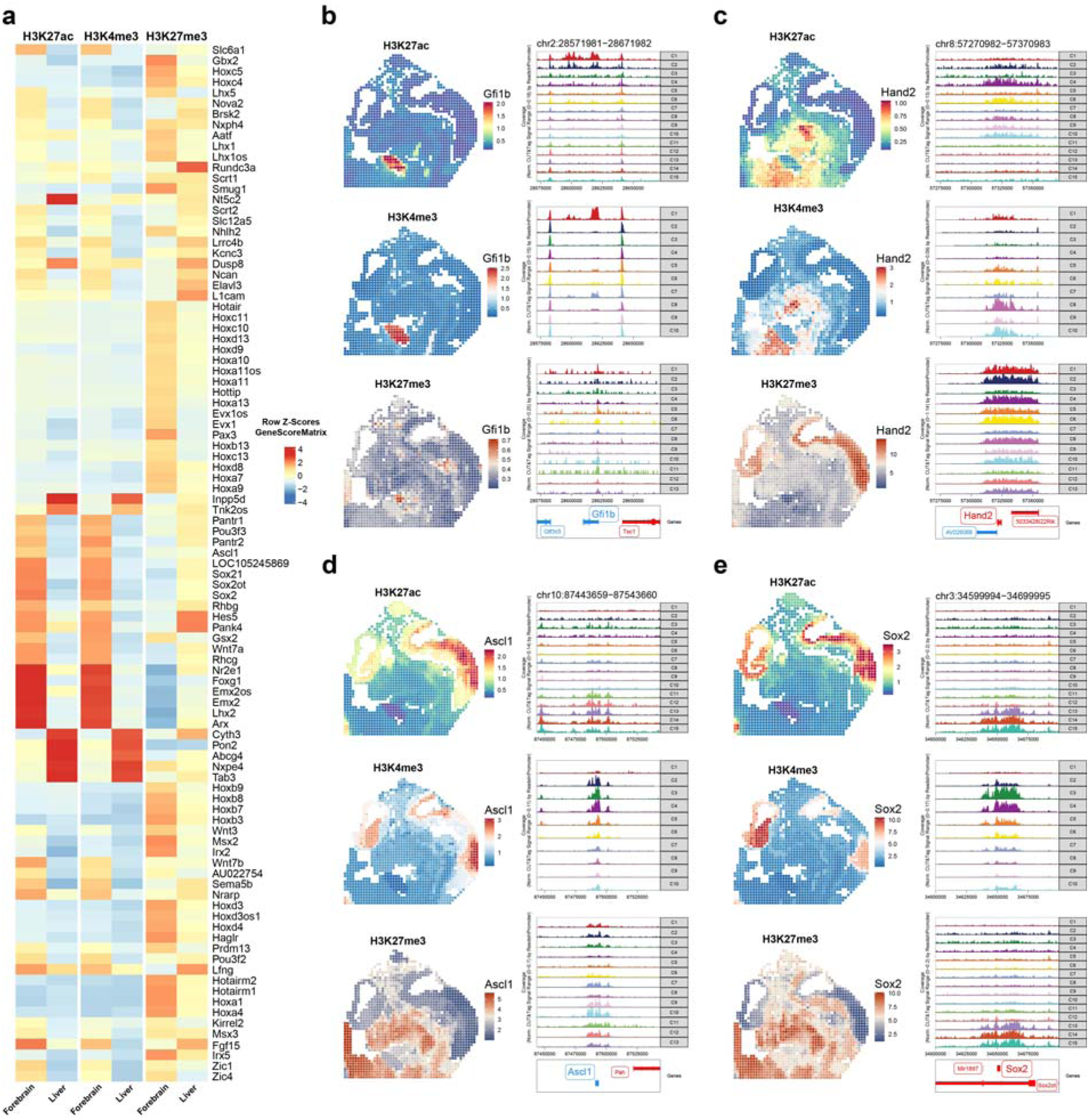
Spatial epigenomic profiling of tissue-specific gene regulation across three histone modifications. **a**, Heatmap illustrating histone modification signals for liver and forebrain marker genes across H3K27ac, H3K4me3 and H3K27me3, highlighting region-specific regulatory signatures. **b**-**e**, Spatial localization maps (left) and genome browser tracks (right) for representative genes, showing tissue-specific chromatin enrichment patterns and corresponding genomic profiles under the three histone marks.

**Extended data Fig. 4.**
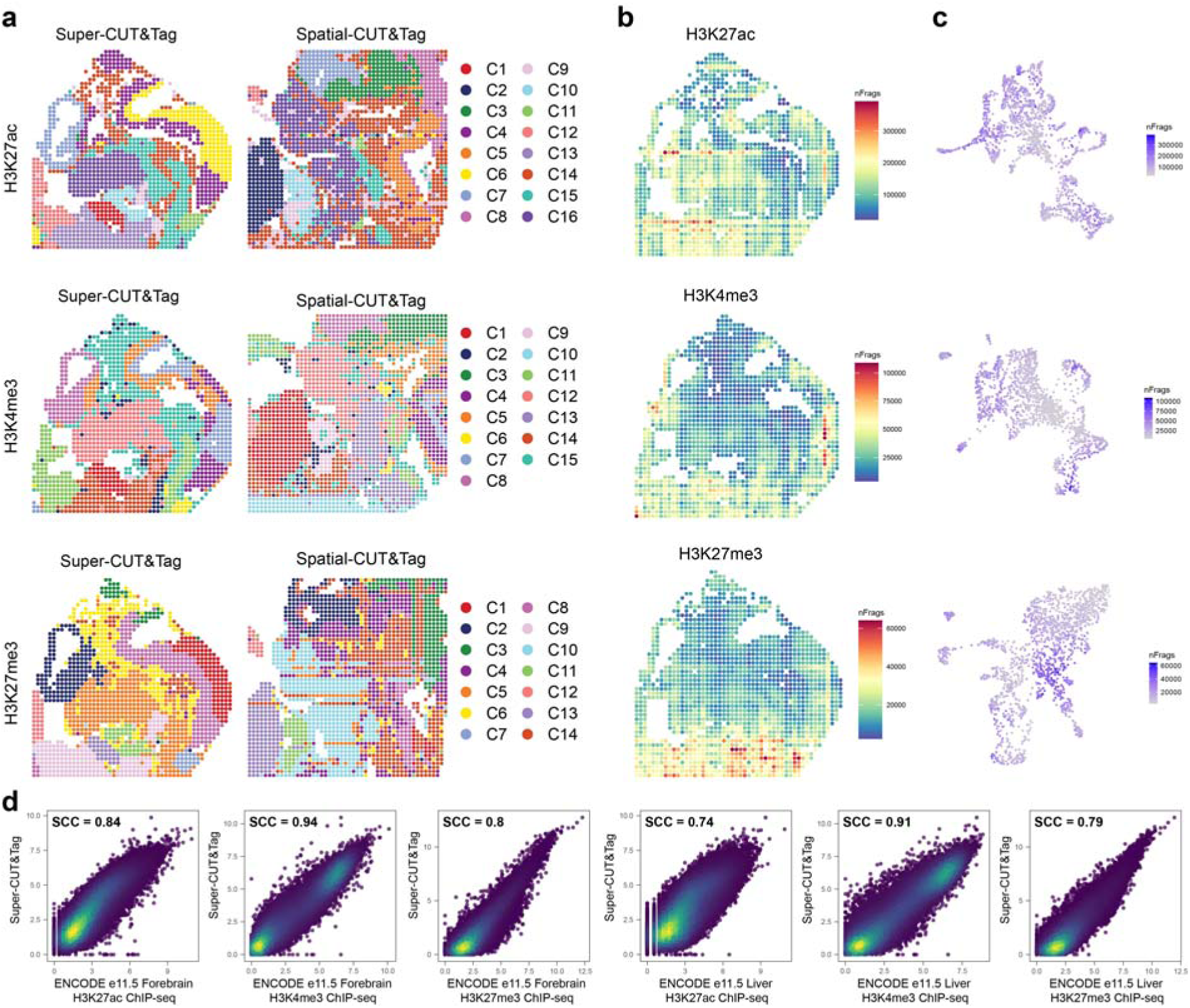
Quality control of e11.5 embryos. **a**, Spatial mapping of clusters from integrated UMAP analysis of Super-CUT&Tag (e11.5) and Spatial-CUT&Tag (e11) chromatin profiles across H3K27ac, H3K4me3 and H3K27me3. **b**, Spatial heatmaps showing unique fragment counts per spot for each histone mark. **c**, UMAP projections of unsupervised clustering results per histone modification, with spot intensity reflecting unique fragment numbers. **d**, Correlation of Super-CUT&Tag signals with ENCODE forebrain and liver datasets across the three histone marks. Spearman correlation coefficients (Spearman’s ρ) indicate concordance between Super-CUT&Tag profiles and ENCODE references.

**Extended data Fig. 5.**
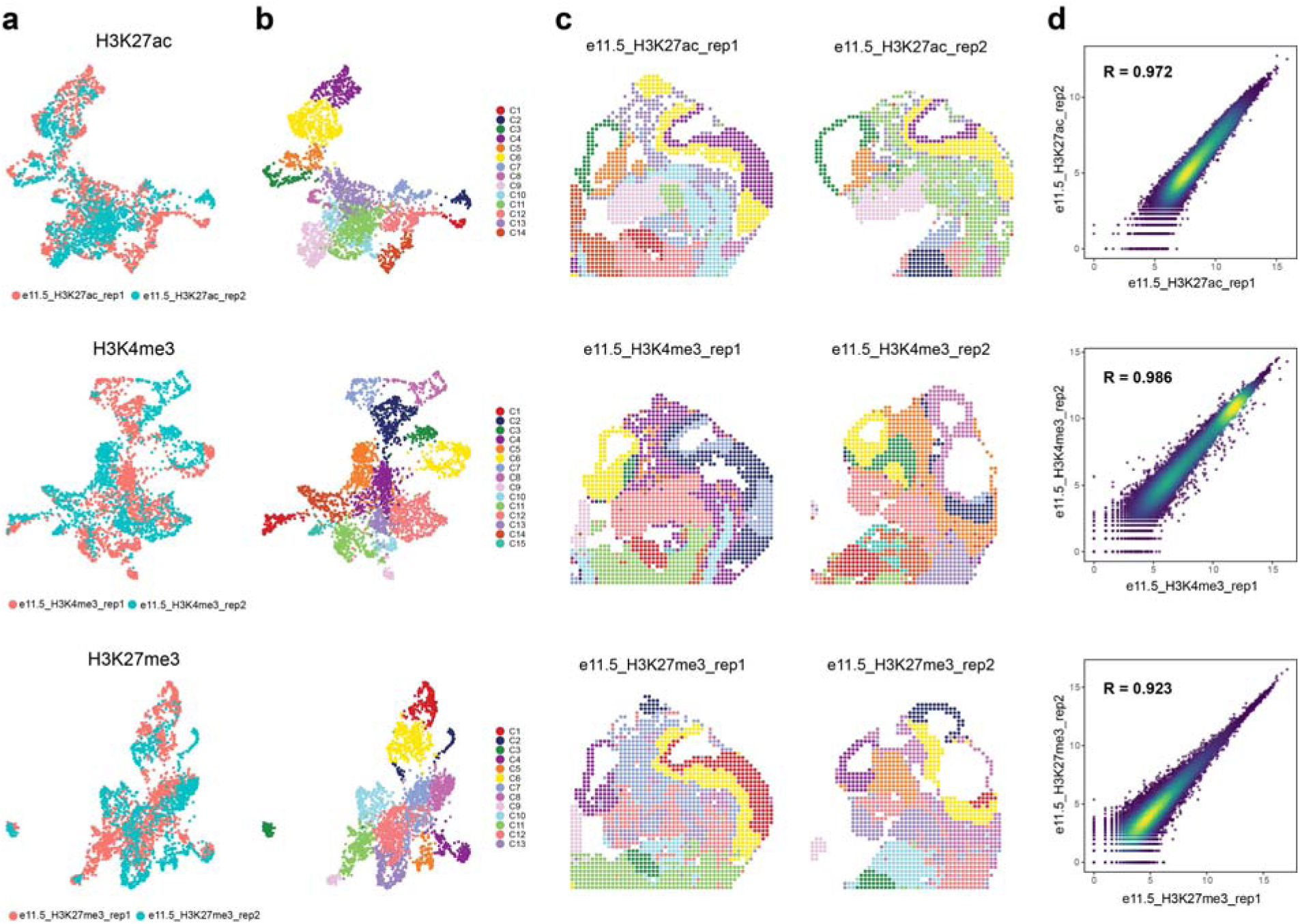
Reproducibility of Super-CUT&Tag. **a,** UMAP projections of Super-CUT&Tag profiles from replicate 1 (red) and replicate 2 (blue). Each dot represents a spatially-resolved spot. The high degree of overlap indicates reproducible spatial epigenomic patterns across replicates. **b,** UMAP projections of the same datasets as in **(a)**, with colors representing unsupervised clusters. Each cluster corresponds to a group of spatial regions with similar chromatin modification profiles. **c,** Spatial maps showing the tissue distribution of clusters identified in **(b)**, respectively. **d,** Spearman correlation coefficient between replicate samples for each histone modification. Each dot represents a spatial bin, with fragment counts compared between replicates. High correlation values indicate strong reproducibility of Super-CUT&Tag profiles across experiments. Each panel **(a-d)** contains three subpanels arranged from top to bottom, corresponding to the histone modifications H3K27ac, H3K4me3 and H3K27me3, respectively.

**Extended data Fig. 6.**
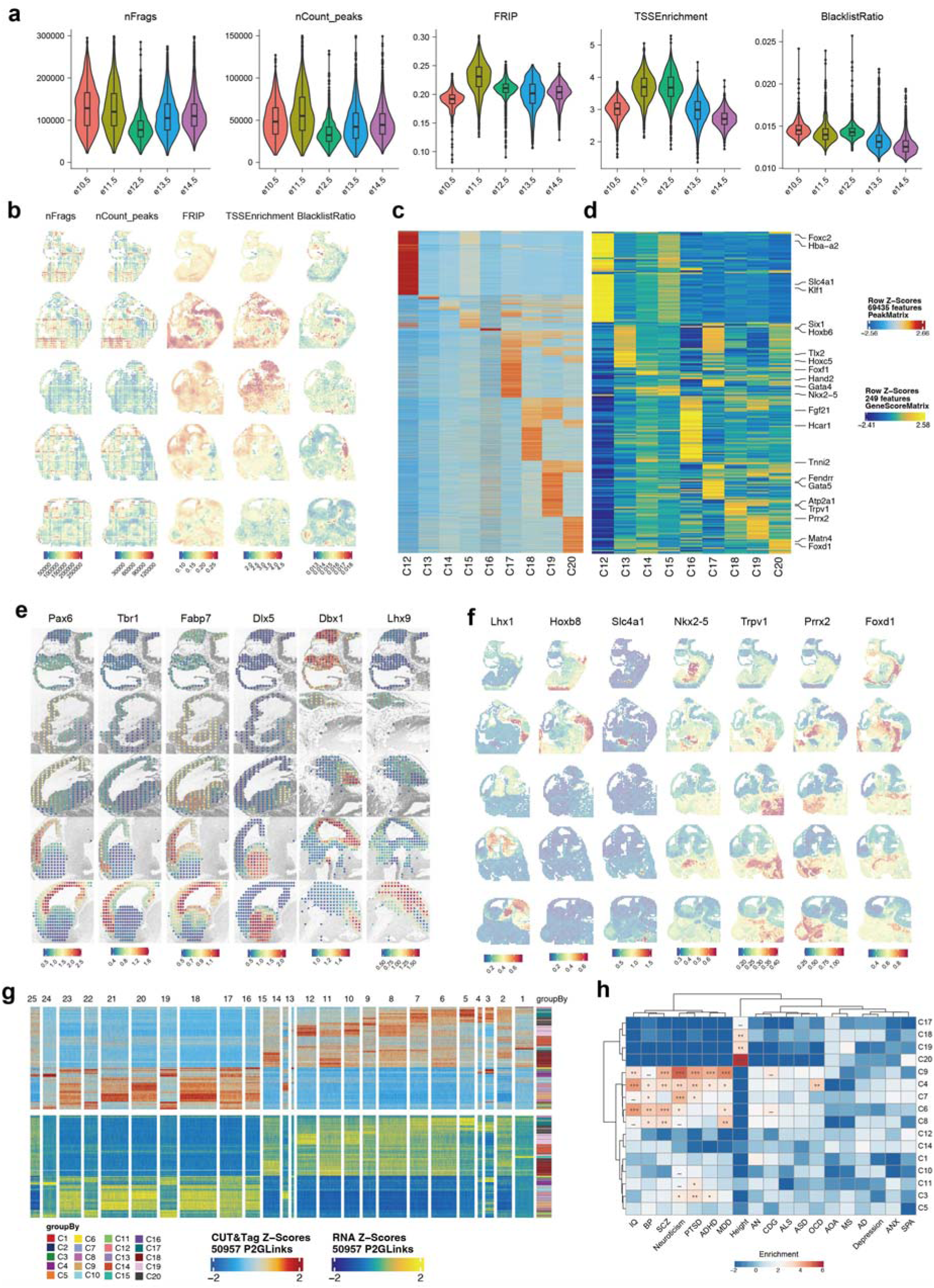
Spatial landscape of H3K27ac modification during mouse embryo development. **a,** Data quality of Super-CUT&Tag on e10.5-e14.5 sections. **b,** Spatial visualization of data qualities. **c-d,** Heatmap of marker peaks (**c**) and marker gene scores (**d**) of clusters C12-C20. **e,** Spatial visualization of H3K27ac modification of marker genes distribution in mouse brain. **f,** Spatial visualization of tissue distribution of marker gene scores. **g,** Heatmap showing H3K27ac peaks and gene expression of 50,927 significant peak-gene linkages. **h**, Human orthologues of H3K27ac peaks exhibit clustered enrichment of noncoding risk variants associated with neurological diseases and traits.

**Extended data Fig. 7.**
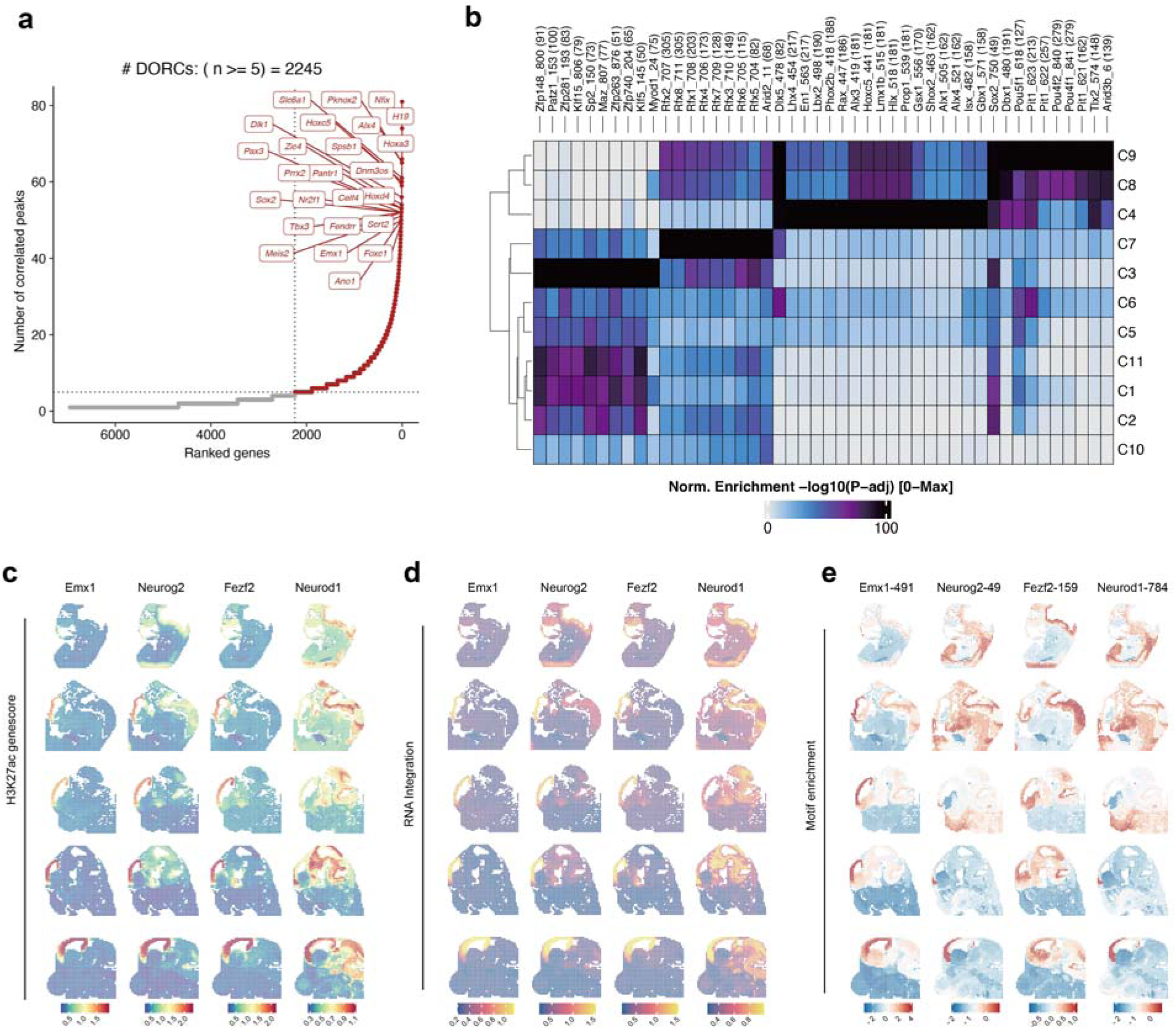
Candidate TF regulators in pallium developments. **a,** Number of significantly correlated peaks per gene. **b**, Motif enrichment of marker peaks in clusters C1-C11. **c-e,** Spatial visualization of H3K27ac gene score (**c**), integrated RNA expression (**d**) and motif enrichment (**e**) for candidate TFs.

**Extended data Fig. 8.**
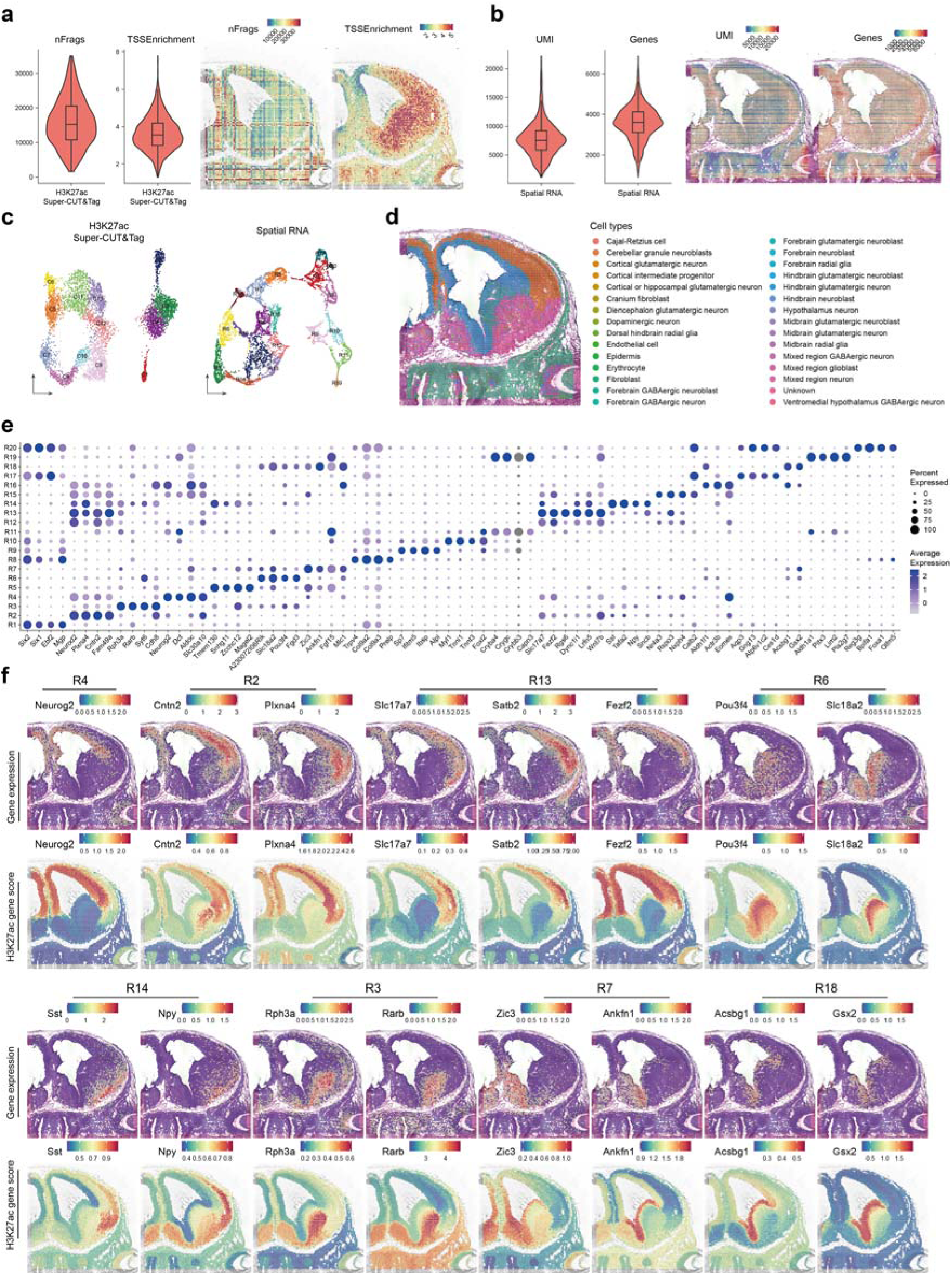
Spatial H3K27ac modification and spatial transcriptome in the e14.5 mouse forebrain. **a-b,** Quality control of H3K27ac Super-CUT&Tag (**a**) and spatial transcriptome (**b**). **c**, UMAP visualization of unsupervised clustering of H3K27ac Super-CUT&Tag data and spatial transcriptomics data. **d**, Spatial distribution of transferred labels from stereo-seq data. **e**, Maker genes of each cluster of spatial transcriptome data. **f**, Spatial expression pattern (top) and H3K27ac gene score (bottom) of marker genes.

**Extended data Fig. 9.**
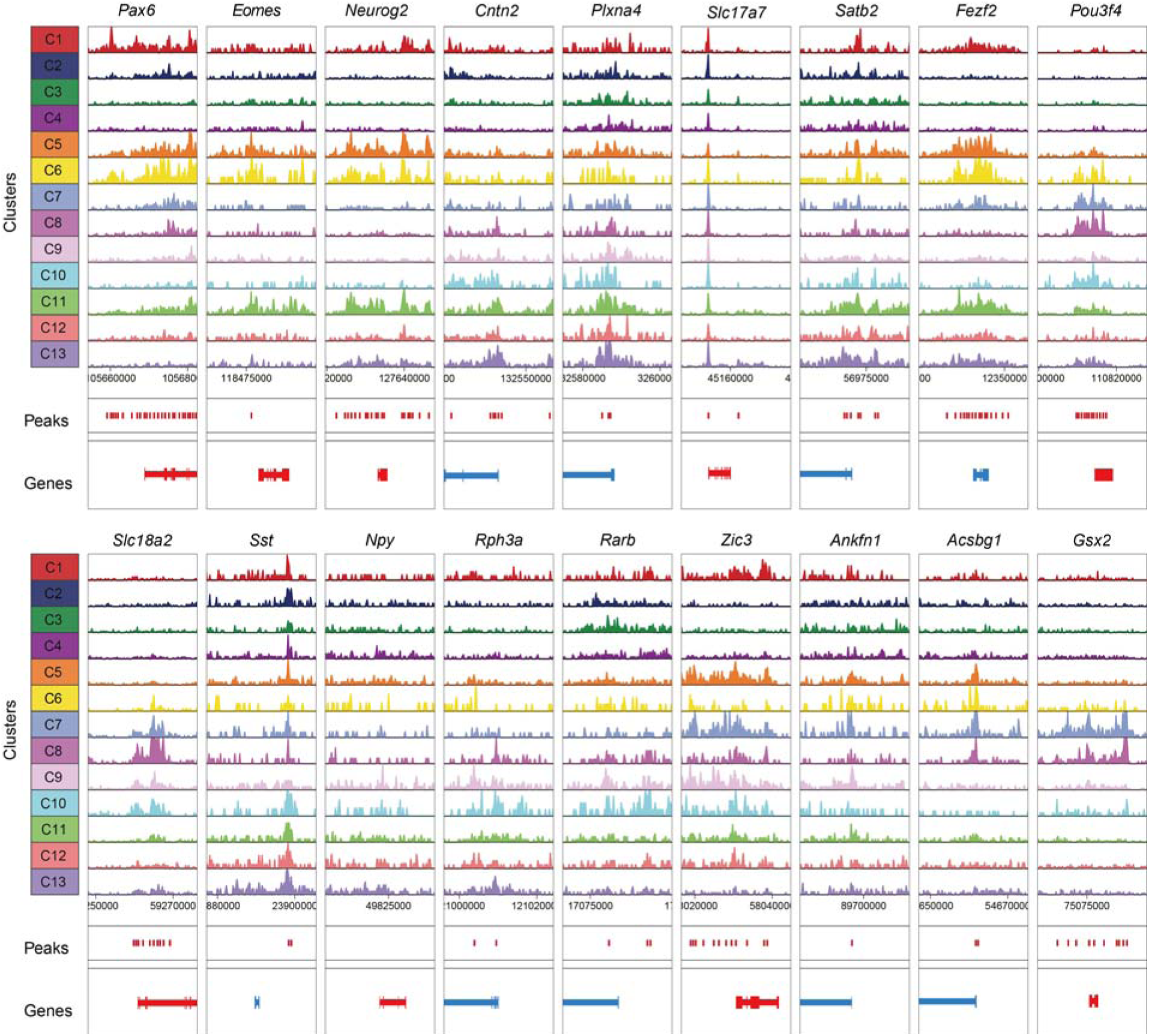
Quality control of H3K27ac Super-CUT&Tag data of forebrain. Genome browser tracks of H3K27ac modification of marker genes across clusters.

**Extended data Fig. 10.**
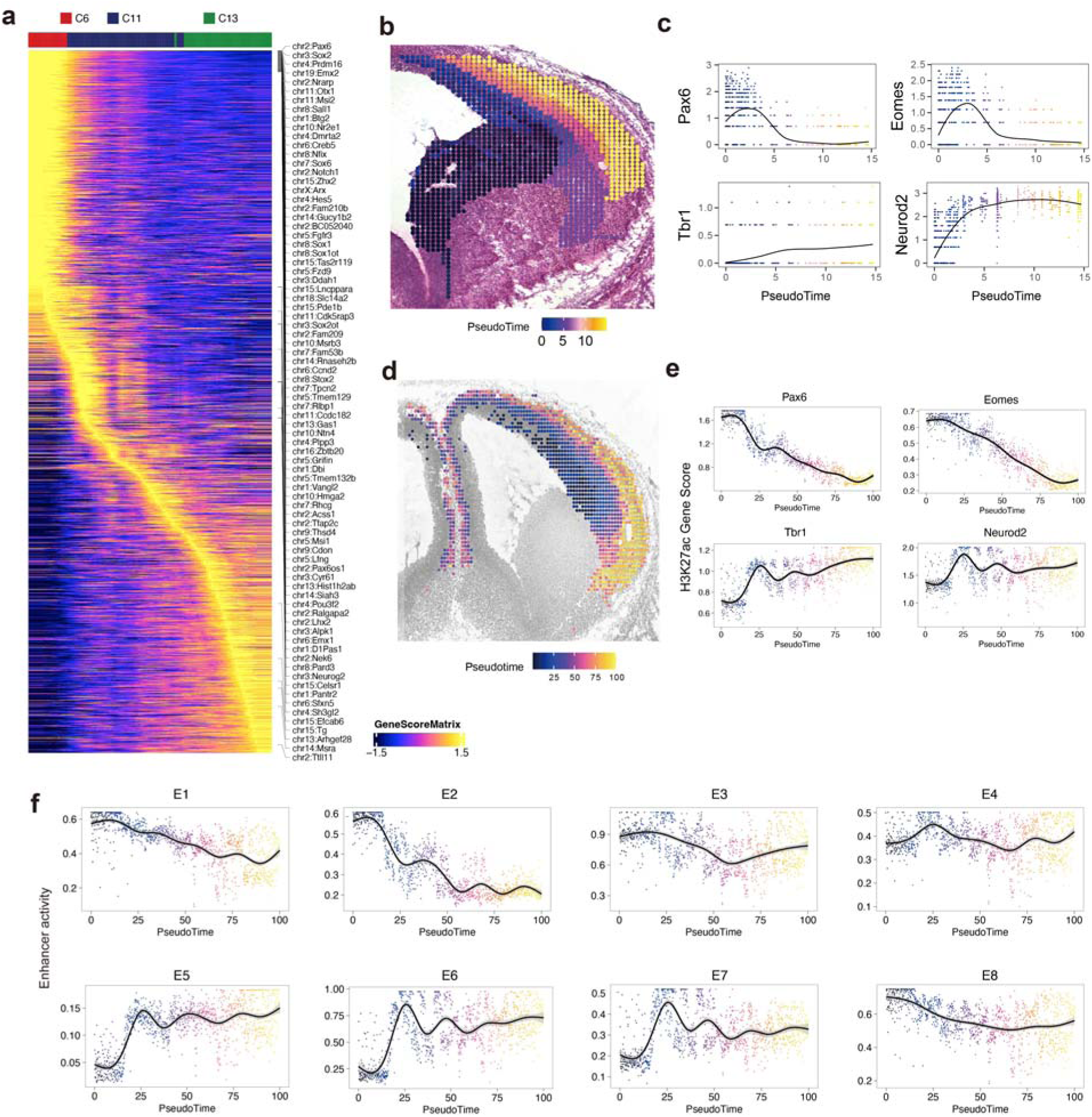
Trajectory analysis of corticogenesis. **a**, Heatmap of H3K27ac Gene Score along pseudotime. **b**, Spatial mapping of the trajectory of corticogenesis of spatial transcriptome data. **c**, Gene expression of *Pax6*, *Eomes*, *Tbr1* and *Neurod2* in the pseudotime axis. **d**, Spatial mapping of the trajectory of corticogenesis of H3K27ac Super-CUT&Tag data. **e**, H3K27ac Gene Score of *Pax6*, *Eomes*, *Tbr1* and *Neurod2* in the pseudotime axis. **f**, Enhancers activity in the pseudotime axis.

