## Supplementary Information for "Super-CUT&Tag: A Sensitive, Spatially Resolved Approach for Epigenomic Profiling in Tissue Sections"

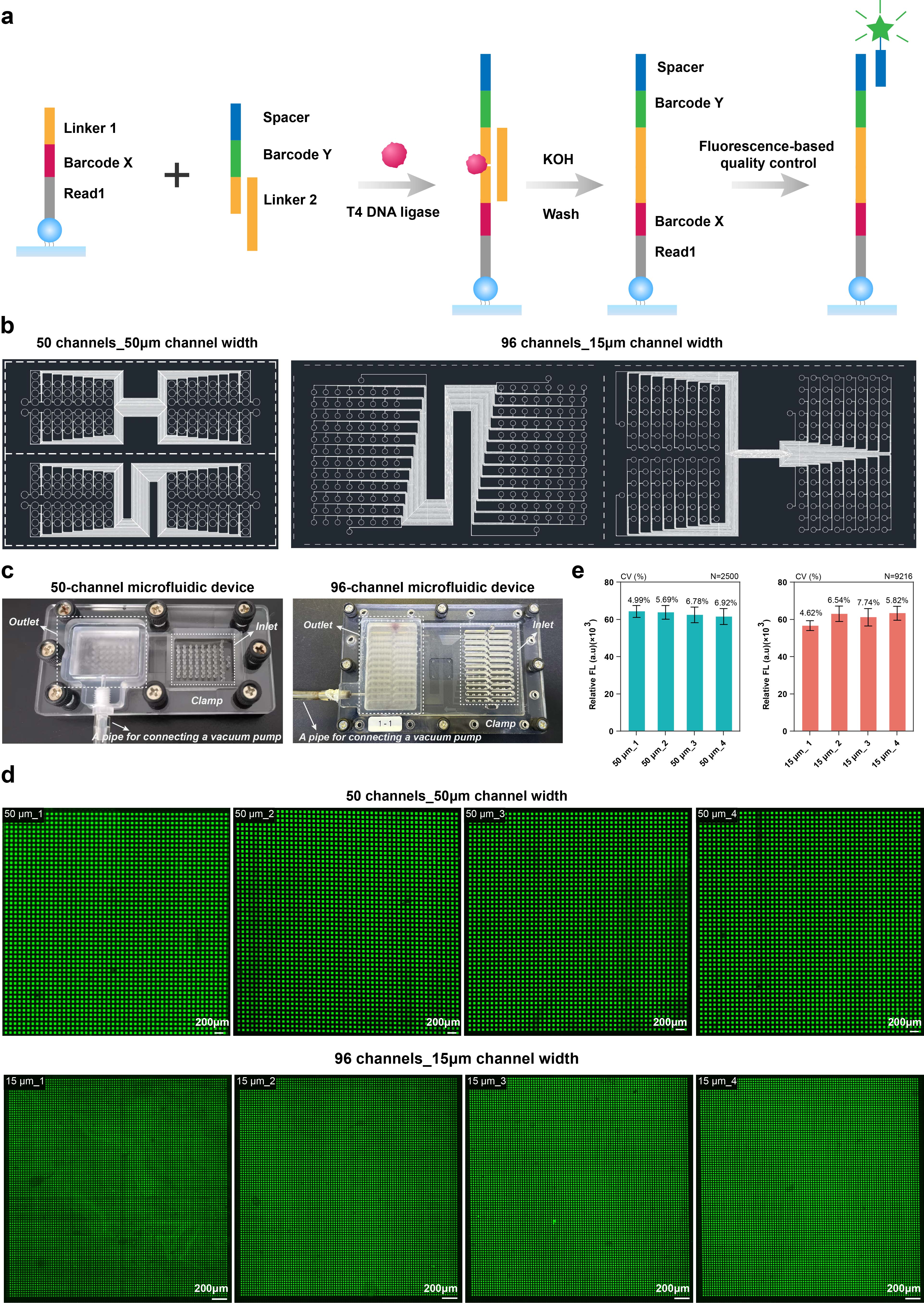
**Supplementary Fig. 1| DNA modification, microfluidic device design, and fluorescence uniformity analysis of three-dimensional (3D) dendrimeric DNA arrays.** **a,** Schematic representation of the DNA ligation reaction and fluorescence-based quality control for barcoding on the 3D dendrimeric slide. **b,** AutoCAD representations of polydimethylsiloxane (PDMS) chips with different channel numbers and widths. Left: 50 channels of 50 μm in width; right: 96 channels of 15 μm in width. **c,** Microfluidic device used for Super-CUT&Tag, featuring a 50-channel design (top) and a 96-channel design (bottom). The PDMS chip was mounted on a slide and clamped with acrylic plates and screws. DNA barcode reagents were loaded into the inlets and drawn through the channels via vacuum applied to the outlet caps. **d,** Fluorescence images of 3D dendrimeric DNA arrays with different spot sizes. The top row shows 50 μm spots, and the bottom row shows 15 μm spots. Scale bar, 200 μm. e, Quantification of fluorescence intensity uniformity across slides. Left: mean fluorescence intensity ± s.d. of 50 μm spots from four slides (N = 2,500 spots per slide). Right: mean fluorescence intensity ± s.d. of 15 μm spots from four slides (N = 9,216 spots per slide). The corresponding coefficients of variation (CV) are indicated.

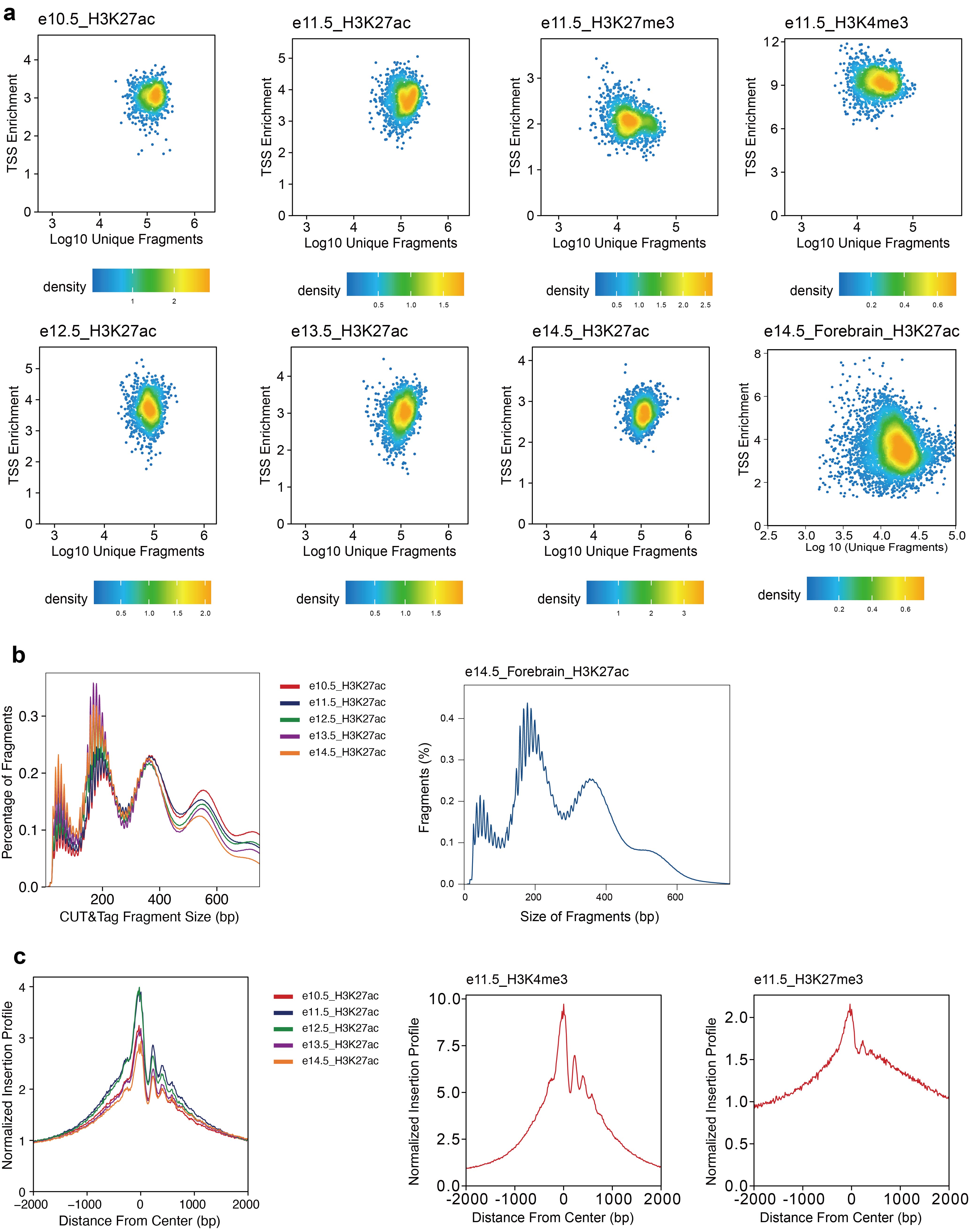

**Supplementary Fig. 2| Quality control of mouse embryo data. a**, Density plots show the log10(unique nuclear fragments) vs TSS enrichment score. **b-c**, Fragment size distributions (**b**) and TSS enrichment profiles (**c**) of each data.

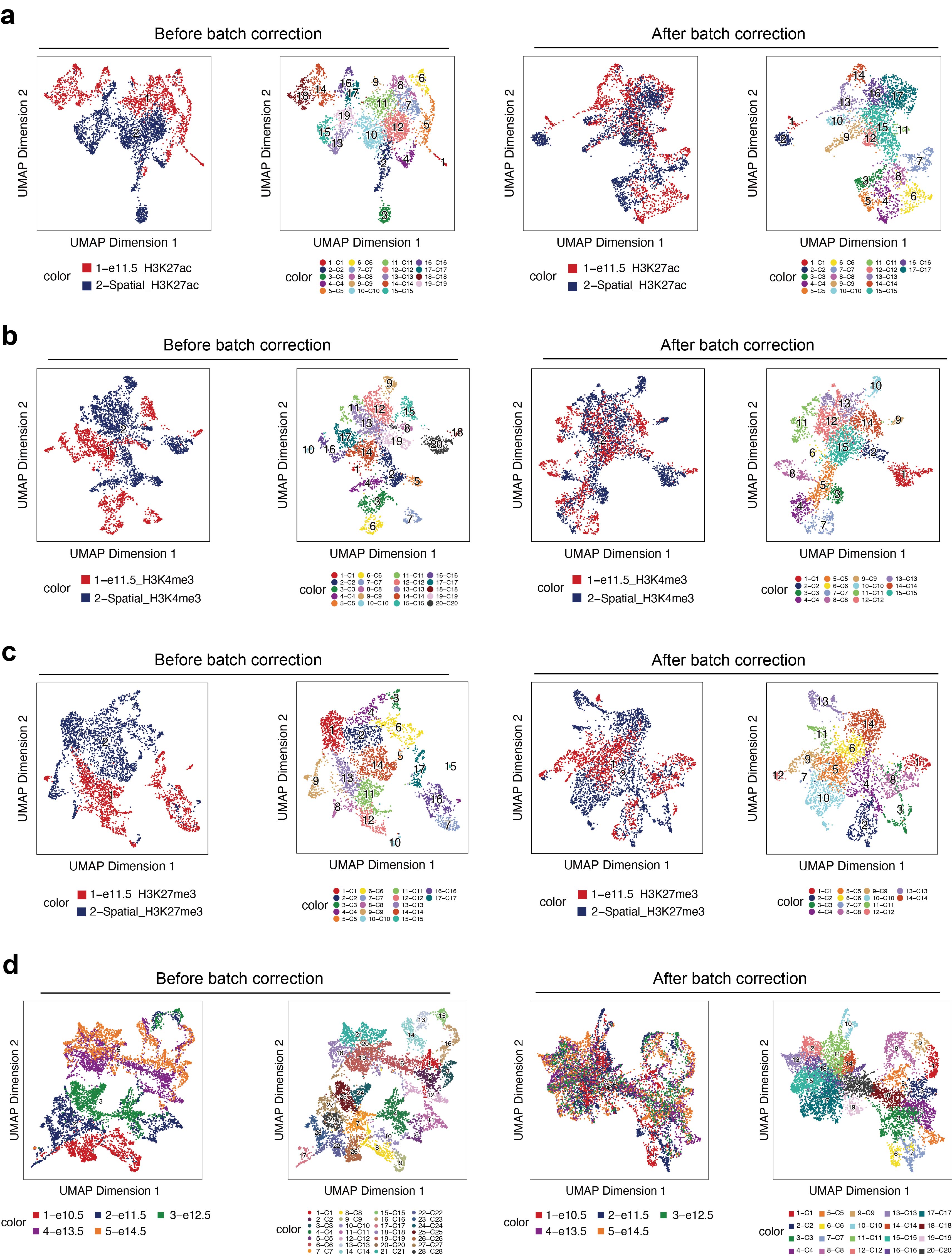

**Supplementary Fig. 3| Batch effects between different spatial epigenomic datasets. a-c**, Comparison of batch effects between Super-CUT&Tag and Spatial-CUT&Tag across three histone modifications. **d**, Batch effects within Super-CUT&Tag datasets across different embryonic developmental stages.

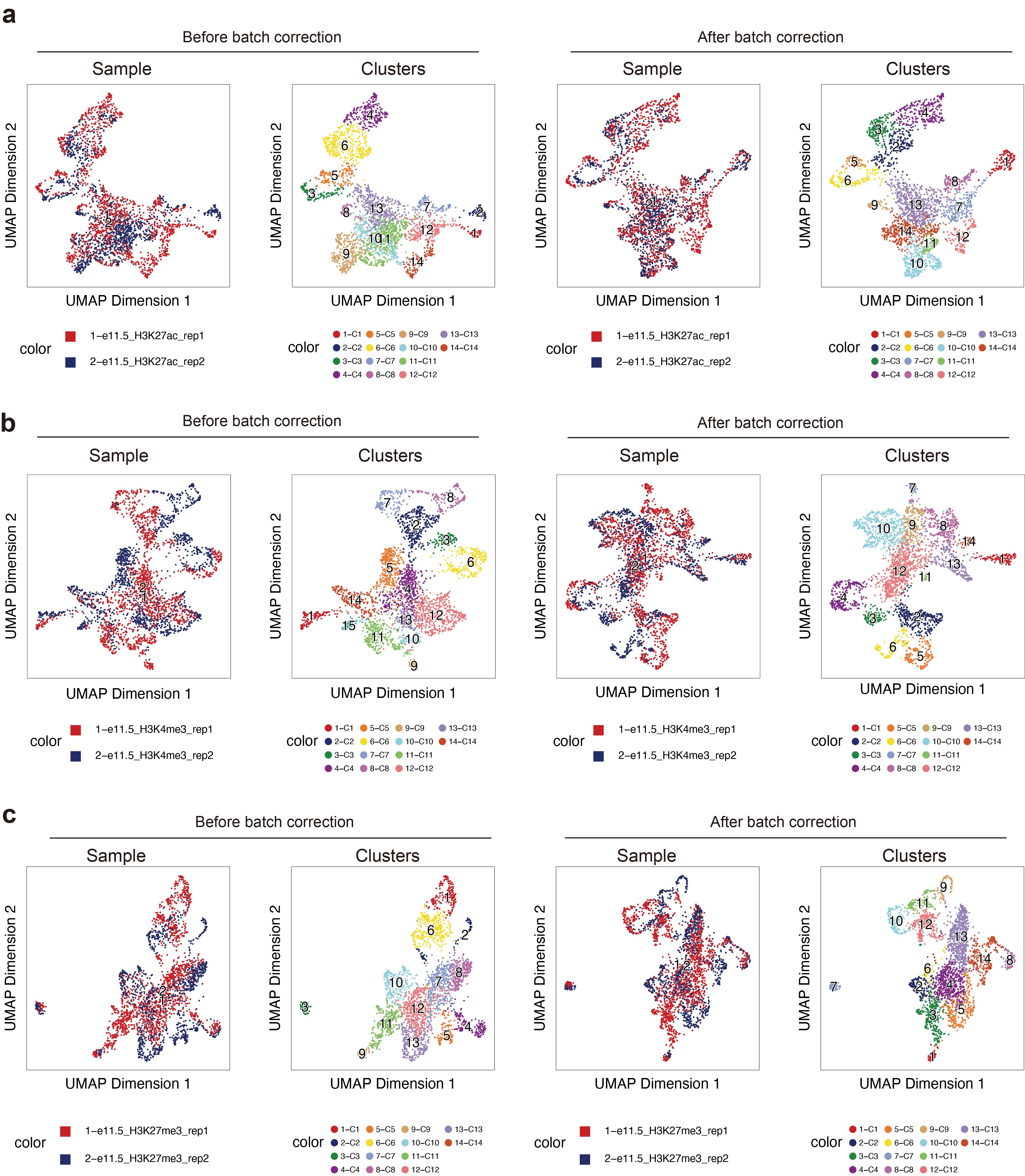

**Supplementary Fig. 4| Batch effects among Super-CUT&Tag replicates. a-c**, Assessment of batch effects within Super-CUT&Tag datasets for three histone modifications across biological replicates.

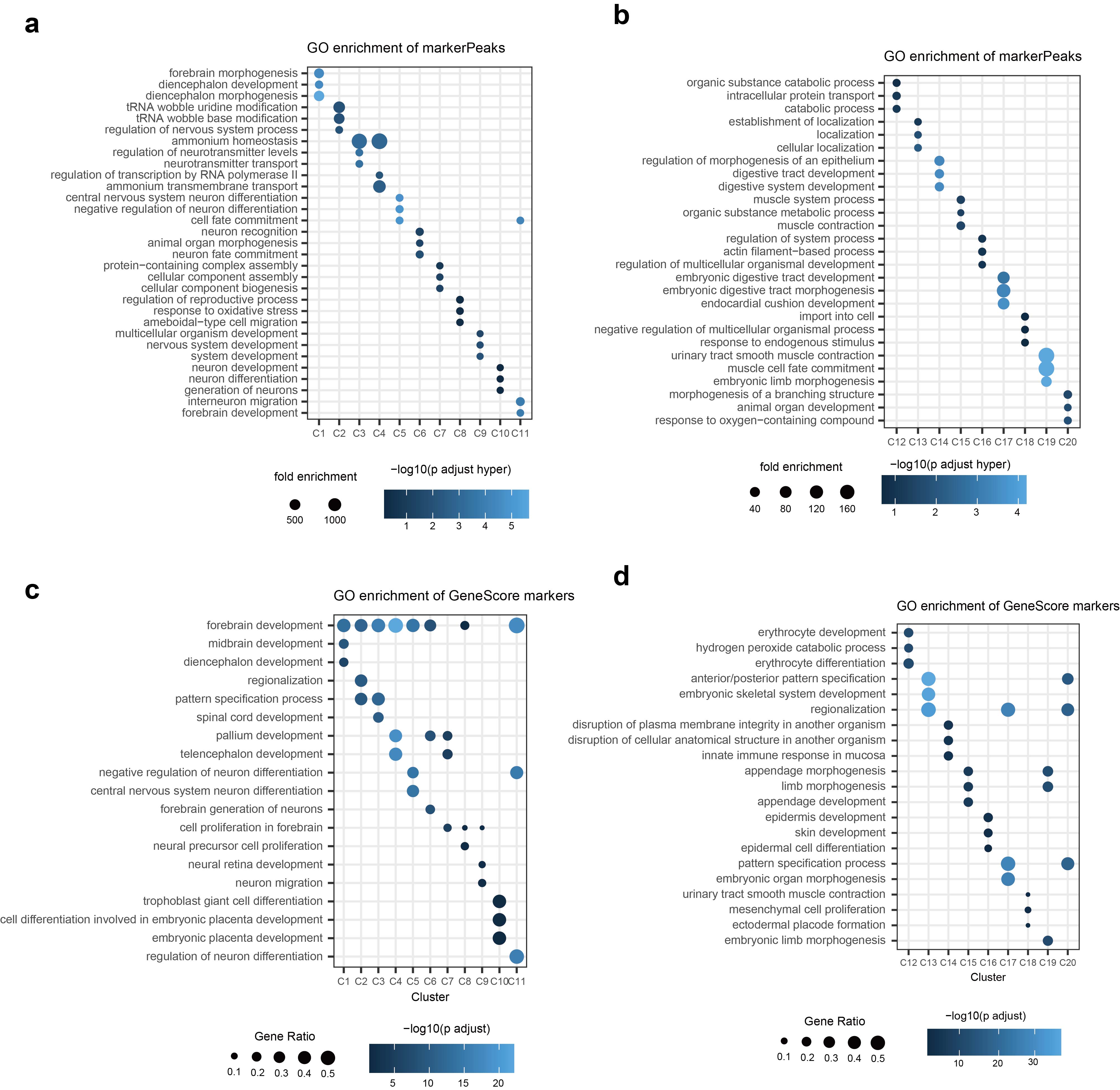

**Supplementary Fig. 5| Function enrichment** **of marker peaks and marker gene scores of data in Fig2. a-b,** GO enrichment analysis of top marker peaks in C1-C11 (**a**) and C12-C20 (**b**). **c-d,** GO enrichment analysis of top marker gene scores in C1-C11 (**c**) and C12-C20 (**d**).

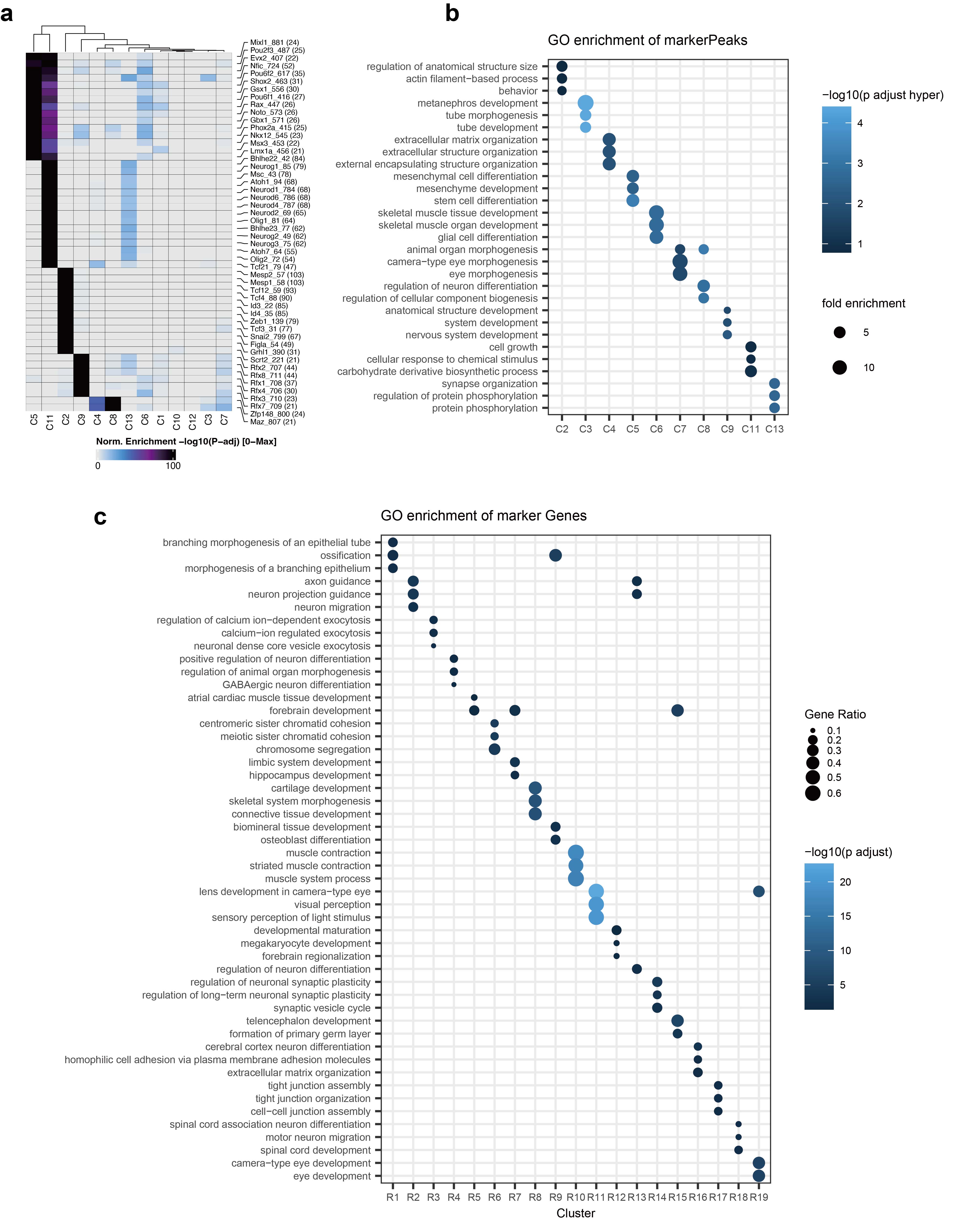
**Supplementary Fig. 6| Function enrichment** **of clusters of data in Fig4. a**, Motif enrichment of H3K27ac Super-CUT&Tag clusters. **b**, GREAT analysis of marker peaks of H3K27ac Super-CUT&Tag clusters. **c**, GO analysis of marker genes of spatial transcriptome clusters of mouse forebrain.

### Supplementary Table 1: Barcode X Sequences

| Barcode X1 | CTACACGACGCTCTTCCGATCTATAATGTCAGGCCAGAGCATTCG |
| --- | --- |
| Barcode X2 | CTACACGACGCTCTTCCGATCTATGCCTAAAGGCCAGAGCATTCG |
| Barcode X3 | CTACACGACGCTCTTCCGATCTAGTGGTCAAGGCCAGAGCATTCG |
| Barcode X4 | CTACACGACGCTCTTCCGATCTACCACTGTAGGCCAGAGCATTCG |
| Barcode X5 | CTACACGACGCTCTTCCGATCTACATTGGCAGGCCAGAGCATTCG |
| Barcode X6 | CTACACGACGCTCTTCCGATCTCCACTTACAGGCCAGAGCATTCG |
| Barcode X7 | CTACACGACGCTCTTCCGATCTCATCAAGTAGGCCAGAGCATTCG |
| Barcode X8 | CTACACGACGCTCTTCCGATCTACAAGCTAAGGCCAGAGCATTCG |
| Barcode X9 | CTACACGACGCTCTTCCGATCTAGTACAAGAGGCCAGAGCATTCG |
| Barcode X10 | CTACACGACGCTCTTCCGATCTAACAACCAAGGCCAGAGCATTCG |
| Barcode X11 | CTACACGACGCTCTTCCGATCTAACCGAGAAGGCCAGAGCATTCG |
| Barcode X12 | CTACACGACGCTCTTCCGATCTAACGCTTAAGGCCAGAGCATTCG |
| Barcode X13 | CTACACGACGCTCTTCCGATCTGAATGCAGAGGCCAGAGCATTCG |
| Barcode X14 | CTACACGACGCTCTTCCGATCTAAGGTACAAGGCCAGAGCATTCG |
| Barcode X15 | CTACACGACGCTCTTCCGATCTACACAGAAAGGCCAGAGCATTCG |
| Barcode X16 | CTACACGACGCTCTTCCGATCTACAGCAGAAGGCCAGAGCATTCG |
| Barcode X17 | CTACACGACGCTCTTCCGATCTACCTCCAAAGGCCAGAGCATTCG |
| Barcode X18 | CTACACGACGCTCTTCCGATCTACGTATCAAGGCCAGAGCATTCG |
| Barcode X19 | CTACACGACGCTCTTCCGATCTACTATGCAAGGCCAGAGCATTCG |
| Barcode X20 | CTACACGACGCTCTTCCGATCTAGAGTCAAAGGCCAGAGCATTCG |
| Barcode X21 | CTACACGACGCTCTTCCGATCTAACGTGATAGGCCAGAGCATTCG |
| Barcode X22 | CTACACGACGCTCTTCCGATCTAGCAGGAAAGGCCAGAGCATTCG |
| Barcode X23 | CTACACGACGCTCTTCCGATCTAGTCACTAAGGCCAGAGCATTCG |
| Barcode X24 | CTACACGACGCTCTTCCGATCTATCCTGTAAGGCCAGAGCATTCG |
| Barcode X25 | CTACACGACGCTCTTCCGATCTGCCAAGACAGGCCAGAGCATTCG |
| Barcode X26 | CTACACGACGCTCTTCCGATCTCAACCACAAGGCCAGAGCATTCG |
| Barcode X27 | CTACACGACGCTCTTCCGATCTGACTAGTAAGGCCAGAGCATTCG |
| Barcode X28 | CTACACGACGCTCTTCCGATCTCAATGGAAAGGCCAGAGCATTCG |
| Barcode X29 | CTACACGACGCTCTTCCGATCTCACTTCGAAGGCCAGAGCATTCG |
| Barcode X30 | CTACACGACGCTCTTCCGATCTCAGCGTTAAGGCCAGAGCATTCG |
| Barcode X31 | CTACACGACGCTCTTCCGATCTCATACCAAAGGCCAGAGCATTCG |
| Barcode X32 | CTACACGACGCTCTTCCGATCTCCAGTTCAAGGCCAGAGCATTCG |
| Barcode X33 | CTACACGACGCTCTTCCGATCTCCGAAGTAAGGCCAGAGCATTCG |
| Barcode X34 | CTACACGACGCTCTTCCGATCTCGAACTTAAGGCCAGAGCATTCG |
| Barcode X35 | CTACACGACGCTCTTCCGATCTCGCATACAAGGCCAGAGCATTCG |
| Barcode X36 | CTACACGACGCTCTTCCGATCTCTCAATGAAGGCCAGAGCATTCG |
| Barcode X37 | CTACACGACGCTCTTCCGATCTCTGGCATAAGGCCAGAGCATTCG |
| Barcode X38 | CTACACGACGCTCTTCCGATCTGAATCTGAAGGCCAGAGCATTCG |
| Barcode X39 | CTACACGACGCTCTTCCGATCTCAAGACTAAGGCCAGAGCATTCG |
| Barcode X40 | CTACACGACGCTCTTCCGATCTGAGCTGAAAGGCCAGAGCATTCG |
| Barcode X41 | CTACACGACGCTCTTCCGATCTGATAGACAAGGCCAGAGCATTCG |
| Barcode X42 | CTACACGACGCTCTTCCGATCTGCCACATAAGGCCAGAGCATTCG |
| Barcode X43 | CTACACGACGCTCTTCCGATCTGCGAGTAAAGGCCAGAGCATTCG |
| Barcode X44 | CTACACGACGCTCTTCCGATCTGCTAACGAAGGCCAGAGCATTCG |
| Barcode X45 | CTACACGACGCTCTTCCGATCTGGAGAACAAGGCCAGAGCATTCG |
| Barcode X46 | CTACACGACGCTCTTCCGATCTGTACGCAAAGGCCAGAGCATTCG |
| Barcode X47 | CTACACGACGCTCTTCCGATCTGTCGTAGAAGGCCAGAGCATTCG |
| Barcode X48 | CTACACGACGCTCTTCCGATCTGTCTGTCAAGGCCAGAGCATTCG |
| Barcode X49 | CTACACGACGCTCTTCCGATCTGTGTTCTAAGGCCAGAGCATTCG |
| Barcode X50 | CTACACGACGCTCTTCCGATCTTAGGATGAAGGCCAGAGCATTCG |
| Barcode X51 | CTACACGACGCTCTTCCGATCTAAACATCGAGGCCAGAGCATTCG |
| Barcode X52 | CTACACGACGCTCTTCCGATCTCGCTGATCAGGCCAGAGCATTCG |
| Barcode X53 | CTACACGACGCTCTTCCGATCTCTGTAGCCAGGCCAGAGCATTCG |
| Barcode X54 | CTACACGACGCTCTTCCGATCTACGCTCGAAGGCCAGAGCATTCG |
| Barcode X55 | CTACACGACGCTCTTCCGATCTCCGTGAGAAGGCCAGAGCATTCG |
| Barcode X56 | CTACACGACGCTCTTCCGATCTCCTCCTGAAGGCCAGAGCATTCG |
| Barcode X57 | CTACACGACGCTCTTCCGATCTCGACTGGAAGGCCAGAGCATTCG |
| Barcode X58 | CTACACGACGCTCTTCCGATCTCTGAGCCAAGGCCAGAGCATTCG |
| Barcode X59 | CTACACGACGCTCTTCCGATCTGCTCGGTAAGGCCAGAGCATTCG |
| Barcode X60 | CTACACGACGCTCTTCCGATCTGGTGCGAAAGGCCAGAGCATTCG |
| Barcode X61 | CTACACGACGCTCTTCCGATCTTATCAGCAAGGCCAGAGCATTCG |
| Barcode X62 | CTACACGACGCTCTTCCGATCTTCCGTCTAAGGCCAGAGCATTCG |
| Barcode X63 | CTACACGACGCTCTTCCGATCTTCTTCACAAGGCCAGAGCATTCG |
| Barcode X64 | CTACACGACGCTCTTCCGATCTTGAAGAGAAGGCCAGAGCATTCG |
| Barcode X65 | CTACACGACGCTCTTCCGATCTTGGAACAAAGGCCAGAGCATTCG |
| Barcode X66 | CTACACGACGCTCTTCCGATCTTGGCTTCAAGGCCAGAGCATTCG |
| Barcode X67 | CTACACGACGCTCTTCCGATCTTGGTGGTAAGGCCAGAGCATTCG |
| Barcode X68 | CTACACGACGCTCTTCCGATCTTTCACGCAAGGCCAGAGCATTCG |
| Barcode X69 | CTACACGACGCTCTTCCGATCTAACTCACCAGGCCAGAGCATTCG |
| Barcode X70 | CTACACGACGCTCTTCCGATCTAAGAGATCAGGCCAGAGCATTCG |
| Barcode X71 | CTACACGACGCTCTTCCGATCTAAGGACACAGGCCAGAGCATTCG |
| Barcode X72 | CTACACGACGCTCTTCCGATCTAATCCGTCAGGCCAGAGCATTCG |
| Barcode X73 | CTACACGACGCTCTTCCGATCTAATGTTGCAGGCCAGAGCATTCG |
| Barcode X74 | CTACACGACGCTCTTCCGATCTACACGACCAGGCCAGAGCATTCG |
| Barcode X75 | CTACACGACGCTCTTCCGATCTACAGATTCAGGCCAGAGCATTCG |
| Barcode X76 | CTACACGACGCTCTTCCGATCTAGATGTACAGGCCAGAGCATTCG |
| Barcode X77 | CTACACGACGCTCTTCCGATCTAGCACCTCAGGCCAGAGCATTCG |
| Barcode X78 | CTACACGACGCTCTTCCGATCTAGCCATGCAGGCCAGAGCATTCG |
| Barcode X79 | CTACACGACGCTCTTCCGATCTAGGCTAACAGGCCAGAGCATTCG |
| Barcode X80 | CTACACGACGCTCTTCCGATCTATAGCGACAGGCCAGAGCATTCG |
| Barcode X81 | CTACACGACGCTCTTCCGATCTATCATTCCAGGCCAGAGCATTCG |
| Barcode X82 | CTACACGACGCTCTTCCGATCTATTGGCTCAGGCCAGAGCATTCG |
| Barcode X83 | CTACACGACGCTCTTCCGATCTCAAGGAGCAGGCCAGAGCATTCG |
| Barcode X84 | CTACACGACGCTCTTCCGATCTCACCTTACAGGCCAGAGCATTCG |
| Barcode X85 | CTACACGACGCTCTTCCGATCTCCATCCTCAGGCCAGAGCATTCG |
| Barcode X86 | CTACACGACGCTCTTCCGATCTCCGACAACAGGCCAGAGCATTCG |
| Barcode X87 | CTACACGACGCTCTTCCGATCTCCTAATCCAGGCCAGAGCATTCG |
| Barcode X88 | CTACACGACGCTCTTCCGATCTCCTCTATCAGGCCAGAGCATTCG |
| Barcode X89 | CTACACGACGCTCTTCCGATCTCGACACACAGGCCAGAGCATTCG |
| Barcode X90 | CTACACGACGCTCTTCCGATCTCGGATTGCAGGCCAGAGCATTCG |
| Barcode X91 | CTACACGACGCTCTTCCGATCTCTAAGGTCAGGCCAGAGCATTCG |
| Barcode X92 | CTACACGACGCTCTTCCGATCTGAACAGGCAGGCCAGAGCATTCG |
| Barcode X93 | CTACACGACGCTCTTCCGATCTGACAGTGCAGGCCAGAGCATTCG |
| Barcode X94 | CTACACGACGCTCTTCCGATCTGAGTTAGCAGGCCAGAGCATTCG |
| Barcode X95 | CTACACGACGCTCTTCCGATCTGATGAATCAGGCCAGAGCATTCG |
| Barcode X96 | CTACACGACGCTCTTCCGATCTATTGAGGAAGGCCAGAGCATTCG |

### Supplementary Table 2: Barcode Y Sequences

| Barcode Y1 | /5Phos/ATCCACGTGCTTGAGAGATCGCACTCGTGTGAAGACAG |
| --- | --- |
| Barcode Y2 | /5Phos/ATCCACGTGCTTGAGATGCCTAACTCGTGTGAAGACAG |
| Barcode Y3 | /5Phos/ATCCACGTGCTTGAGAGTGGTCACTCGTGTGAAGACAG |
| Barcode Y4 | /5Phos/ATCCACGTGCTTGAGACCACTGTCTCGTGTGAAGACAG |
| Barcode Y5 | /5Phos/ATCCACGTGCTTGAGACATTGGCCTCGTGTGAAGACAG |
| Barcode Y6 | /5Phos/ATCCACGTGCTTGAGGCTCTTCACTCGTGTGAAGACAG |
| Barcode Y7 | /5Phos/ATCCACGTGCTTGAGCATCAAGTCTCGTGTGAAGACAG |
| Barcode Y8 | /5Phos/ATCCACGTGCTTGAGACAAGCTACTCGTGTGAAGACAG |
| Barcode Y9 | /5Phos/ATCCACGTGCTTGAGAGTACAAGCTCGTGTGAAGACAG |
| Barcode Y10 | /5Phos/ATCCACGTGCTTGAGAACAACCACTCGTGTGAAGACAG |
| Barcode Y11 | /5Phos/ATCCACGTGCTTGAGAACCGAGACTCGTGTGAAGACAG |
| Barcode Y12 | /5Phos/ATCCACGTGCTTGAGAACGCTTACTCGTGTGAAGACAG |
| Barcode Y13 | /5Phos/ATCCACGTGCTTGAGAAGACGGACTCGTGTGAAGACAG |
| Barcode Y14 | /5Phos/ATCCACGTGCTTGAGAAGGTACACTCGTGTGAAGACAG |
| Barcode Y15 | /5Phos/ATCCACGTGCTTGAGACACAGAACTCGTGTGAAGACAG |
| Barcode Y16 | /5Phos/ATCCACGTGCTTGAGACAGCAGACTCGTGTGAAGACAG |
| Barcode Y17 | /5Phos/ATCCACGTGCTTGAGACCTCCAACTCGTGTGAAGACAG |
| Barcode Y18 | /5Phos/ATCCACGTGCTTGAGACGTATCACTCGTGTGAAGACAG |
| Barcode Y19 | /5Phos/ATCCACGTGCTTGAGACTATGCACTCGTGTGAAGACAG |
| Barcode Y20 | /5Phos/ATCCACGTGCTTGAGAGAGTCAACTCGTGTGAAGACAG |
| Barcode Y21 | /5Phos/ATCCACGTGCTTGAGAACGTGATCTCGTGTGAAGACAG |
| Barcode Y22 | /5Phos/ATCCACGTGCTTGAGAGCAGGAACTCGTGTGAAGACAG |
| Barcode Y23 | /5Phos/ATCCACGTGCTTGAGAGTCACTACTCGTGTGAAGACAG |
| Barcode Y24 | /5Phos/ATCCACGTGCTTGAGATCCTGTACTCGTGTGAAGACAG |
| Barcode Y25 | /5Phos/ATCCACGTGCTTGAGGCCAAGACCTCGTGTGAAGACAG |
| Barcode Y26 | /5Phos/ATCCACGTGCTTGAGCAACCACACTCGTGTGAAGACAG |
| Barcode Y27 | /5Phos/ATCCACGTGCTTGAGGACTAGTACTCGTGTGAAGACAG |
| Barcode Y28 | /5Phos/ATCCACGTGCTTGAGCAATGGAACTCGTGTGAAGACAG |
| Barcode Y29 | /5Phos/ATCCACGTGCTTGAGCACTTCGACTCGTGTGAAGACAG |
| Barcode Y30 | /5Phos/ATCCACGTGCTTGAGCAGCGTTACTCGTGTGAAGACAG |
| Barcode Y31 | /5Phos/ATCCACGTGCTTGAGCATACCAACTCGTGTGAAGACAG |
| Barcode Y32 | /5Phos/ATCCACGTGCTTGAGCCAGTTCACTCGTGTGAAGACAG |
| Barcode Y33 | /5Phos/ATCCACGTGCTTGAGCCGAAGTACTCGTGTGAAGACAG |
| Barcode Y34 | /5Phos/ATCCACGTGCTTGAGCGAACTTACTCGTGTGAAGACAG |
| Barcode Y35 | /5Phos/ATCCACGTGCTTGAGCGCATACACTCGTGTGAAGACAG |
| Barcode Y36 | /5Phos/ATCCACGTGCTTGAGCTCAATGACTCGTGTGAAGACAG |
| Barcode Y37 | /5Phos/ATCCACGTGCTTGAGCTGGCATACTCGTGTGAAGACAG |
| Barcode Y38 | /5Phos/ATCCACGTGCTTGAGGAATCTGACTCGTGTGAAGACAG |
| Barcode Y39 | /5Phos/ATCCACGTGCTTGAGCAAGACTACTCGTGTGAAGACAG |
| Barcode Y40 | /5Phos/ATCCACGTGCTTGAGGAGCTGAACTCGTGTGAAGACAG |
| Barcode Y41 | /5Phos/ATCCACGTGCTTGAGGATAGACACTCGTGTGAAGACAG |
| Barcode Y42 | /5Phos/ATCCACGTGCTTGAGGCCACATACTCGTGTGAAGACAG |
| Barcode Y43 | /5Phos/ATCCACGTGCTTGAGGCGAGTAACTCGTGTGAAGACAG |
| Barcode Y44 | /5Phos/ATCCACGTGCTTGAGGCTAACGACTCGTGTGAAGACAG |
| Barcode Y45 | /5Phos/ATCCACGTGCTTGAGGGAGAACACTCGTGTGAAGACAG |
| Barcode Y46 | /5Phos/ATCCACGTGCTTGAGGTACGCAACTCGTGTGAAGACAG |
| Barcode Y47 | /5Phos/ATCCACGTGCTTGAGGTCGTAGACTCGTGTGAAGACAG |
| Barcode Y48 | /5Phos/ATCCACGTGCTTGAGGTCTGTCACTCGTGTGAAGACAG |
| Barcode Y49 | /5Phos/ATCCACGTGCTTGAGGTGTTCTACTCGTGTGAAGACAG |
| Barcode Y50 | /5Phos/ATCCACGTGCTTGAGTAGGATGACTCGTGTGAAGACAG |
| Barcode Y51 | /5Phos/ATCCACGTGCTTGAGAAACATCGCTCGTGTGAAGACAG |
| Barcode Y52 | /5Phos/ATCCACGTGCTTGAGCGCTGATCCTCGTGTGAAGACAG |
| Barcode Y53 | /5Phos/ATCCACGTGCTTGAGCTGTAGCCCTCGTGTGAAGACAG |
| Barcode Y54 | /5Phos/ATCCACGTGCTTGAGACGCTCGACTCGTGTGAAGACAG |
| Barcode Y55 | /5Phos/ATCCACGTGCTTGAGCCGTGAGACTCGTGTGAAGACAG |
| Barcode Y56 | /5Phos/ATCCACGTGCTTGAGCCTCCTGACTCGTGTGAAGACAG |
| Barcode Y57 | /5Phos/ATCCACGTGCTTGAGCGACTGGACTCGTGTGAAGACAG |
| Barcode Y58 | /5Phos/ATCCACGTGCTTGAGCTGAGCCACTCGTGTGAAGACAG |
| Barcode Y59 | /5Phos/ATCCACGTGCTTGAGGCTCGGTACTCGTGTGAAGACAG |
| Barcode Y60 | /5Phos/ATCCACGTGCTTGAGGGTGCGAACTCGTGTGAAGACAG |
| Barcode Y61 | /5Phos/ATCCACGTGCTTGAGTATCAGCACTCGTGTGAAGACAG |
| Barcode Y62 | /5Phos/ATCCACGTGCTTGAGTCCGTCTACTCGTGTGAAGACAG |
| Barcode Y63 | /5Phos/ATCCACGTGCTTGAGTCTTCACACTCGTGTGAAGACAG |
| Barcode Y64 | /5Phos/ATCCACGTGCTTGAGTGAAGAGACTCGTGTGAAGACAG |
| Barcode Y65 | /5Phos/ATCCACGTGCTTGAGTGGAACAACTCGTGTGAAGACAG |
| Barcode Y66 | /5Phos/ATCCACGTGCTTGAGTGGCTTCACTCGTGTGAAGACAG |
| Barcode Y67 | /5Phos/ATCCACGTGCTTGAGTGGTGGTACTCGTGTGAAGACAG |
| Barcode Y68 | /5Phos/ATCCACGTGCTTGAGTTCACGCACTCGTGTGAAGACAG |
| Barcode Y69 | /5Phos/ATCCACGTGCTTGAGAACTCACCCTCGTGTGAAGACAG |
| Barcode Y70 | /5Phos/ATCCACGTGCTTGAGAAGAGATCCTCGTGTGAAGACAG |
| Barcode Y71 | /5Phos/ATCCACGTGCTTGAGAAGGACACCTCGTGTGAAGACAG |
| Barcode Y72 | /5Phos/ATCCACGTGCTTGAGAATCCGTCCTCGTGTGAAGACAG |
| Barcode Y73 | /5Phos/ATCCACGTGCTTGAGAATGTTGCCTCGTGTGAAGACAG |
| Barcode Y74 | /5Phos/ATCCACGTGCTTGAGACACGACCCTCGTGTGAAGACAG |
| Barcode Y75 | /5Phos/ATCCACGTGCTTGAGACAGATTCCTCGTGTGAAGACAG |
| Barcode Y76 | /5Phos/ATCCACGTGCTTGAGAGATGTACCTCGTGTGAAGACAG |
| Barcode Y77 | /5Phos/ATCCACGTGCTTGAGAGCACCTCCTCGTGTGAAGACAG |
| Barcode Y78 | /5Phos/ATCCACGTGCTTGAGAGCCATGCCTCGTGTGAAGACAG |
| Barcode Y79 | /5Phos/ATCCACGTGCTTGAGAGGCTAACCTCGTGTGAAGACAG |
| Barcode Y80 | /5Phos/ATCCACGTGCTTGAGATAGCGACCTCGTGTGAAGACAG |
| Barcode Y81 | /5Phos/ATCCACGTGCTTGAGATCATTCCCTCGTGTGAAGACAG |
| Barcode Y82 | /5Phos/ATCCACGTGCTTGAGATTGGCTCCTCGTGTGAAGACAG |
| Barcode Y83 | /5Phos/ATCCACGTGCTTGAGCAAGGAGCCTCGTGTGAAGACAG |
| Barcode Y84 | /5Phos/ATCCACGTGCTTGAGCACCTTACCTCGTGTGAAGACAG |
| Barcode Y85 | /5Phos/ATCCACGTGCTTGAGCCATCCTCCTCGTGTGAAGACAG |
| Barcode Y86 | /5Phos/ATCCACGTGCTTGAGCCGACAACCTCGTGTGAAGACAG |
| Barcode Y87 | /5Phos/ATCCACGTGCTTGAGCCTAATCCCTCGTGTGAAGACAG |
| Barcode Y88 | /5Phos/ATCCACGTGCTTGAGCCTCTATCCTCGTGTGAAGACAG |
| Barcode Y89 | /5Phos/ATCCACGTGCTTGAGCGACACACCTCGTGTGAAGACAG |
| Barcode Y90 | /5Phos/ATCCACGTGCTTGAGCGGATTGCCTCGTGTGAAGACAG |
| Barcode Y91 | /5Phos/ATCCACGTGCTTGAGCTAAGGTCCTCGTGTGAAGACAG |
| Barcode Y92 | /5Phos/ATCCACGTGCTTGAGGAACAGGCCTCGTGTGAAGACAG |
| Barcode Y93 | /5Phos/ATCCACGTGCTTGAGGACAGTGCCTCGTGTGAAGACAG |
| Barcode Y94 | /5Phos/ATCCACGTGCTTGAGGAGTTAGCCTCGTGTGAAGACAG |
| Barcode Y95 | /5Phos/ATCCACGTGCTTGAGGATGAATCCTCGTGTGAAGACAG |
| Barcode Y96 | /5Phos/ATCCACGTGCTTGAGATTGAGGACTCGTGTGAAGACAG |

### Supplementary Table 3: DNA oligos used for PCR and preparation of sequencing library

| barcode X | NH2-C6-CTACACGACGCTCTTCCGATCT [barcode X] AGGCCAGAGCATTCG |
| --- | --- |
| barcode Y | /5Phos/ATCCACGTGCTTGAG [barcode Y]CTCGTGTGAAGACAG |
| Ligation linker | CTCAAGCACGTGGATCGAATGCTCTGGCCT |
| Tn5MErev | /5Phos/CTGTCTCTTATACACATCT |
| Tn5ME-A | /5Phos/CAAGTATGCAGCGCGAGATGTGTATAAGAGACAG |
| Tn5ME-B | GTCTCGTGGGCTCGGAGATGTGTATAAGAGACAG |
| Splint oligo | /5Phos/CGCGCTGCATACTTGCTGTCTTCACACGAG |
| Forward primer | CTACACGACGCTCTTCCGATCT |
| Reverse primer | GTCTCGTGGGCTCGGAGA |
| P501 | AATGATACGGCGACCACCGAGATCTACACCTATTAAGACACTCTTTCCCTACACGACGCTCTTCCG |
| N701 | CAAGCAGAAGACGGCATACGAGATTCGCCTTAGTCTCGTGGGCTCGG |
| N702 | CAAGCAGAAGACGGCATACGAGATCTAGTACGGTCTCGTGGGCTCGG |
| N703 | CAAGCAGAAGACGGCATACGAGATTTCTGCCTGTCTCGTGGGCTCGG |
| N704 | CAAGCAGAAGACGGCATACGAGATGCTCAGGAGTCTCGTGGGCTCGG |
| N705 | CAAGCAGAAGACGGCATACGAGATAGGAGTCCGTCTCGTGGGCTCGG |
| N706 | CAAGCAGAAGACGGCATACGAGATCATGCCTAGTCTCGTGGGCTCGG |
| N707 | CAAGCAGAAGACGGCATACGAGATGTAGAGAGGTCTCGTGGGCTCGG |
| N708 | CAAGCAGAAGACGGCATACGAGATCCTCTCTGGTCTCGTGGGCTCGG |

### Supplementary Table 4: Encode ChIP-seq peaks used in this study

| **Tissue** | **H3K4me3** | **H3K27me3** | **H3K27ac** |
| --- | --- | --- | --- |
| hindbrain | ENCFF292XPT | ENCFF641DRR | ENCFF083MLY |
| midbrain | ENCFF098WIS | ENCFF927GUC | ENCFF650WFB |
| limb | ENCFF388OWQ | ENCFF601DCY | ENCFF016BEF |
| forebrain | ENCFF635RVF | ENCFF013NFN | ENCFF759KHX |
| lung | ENCFF082TBM | - | - |
| embryonic facial prominence | ENCFF902KTA | ENCFF533OZG | ENCFF680UPD |
| heart | ENCFF908XCE | ENCFF043FMD | ENCFF236UMU |
| neural tube | ENCFF081YTE | ENCFF986DZV | ENCFF554YRS |
| liver | ENCFF220MRB | ENCFF691RWK | ENCFF042BGY |

### Supplementary Table 5: Public data used in this study

| Factor | CistromeDB | GEO or ENCODE | Biological Source | Citation |
| --- | --- | --- | --- | --- |
| PAX6 | 55368 | [GSM1635069](http://www.ncbi.nlm.nih.gov/geo/query/acc.cgi?acc=GSM1635069) | Tissue: Forebrain | Sun J, et al.^1^ |
| CTCF | 64188 | [ENCSR677HXC_1](https://www.encodeproject.org/experiments/ENCSR677HXC) | Tissue: Forebrain | Davis CA, et al.^2^ |
| NEUROG2 | 70732 | [GSM1553880](http://www.ncbi.nlm.nih.gov/geo/query/acc.cgi?acc=GSM1553880) | Cell Type: Cortex | Sessa A, et al.^3^ |
| TBR2 | 70733 | [GSM1553879](http://www.ncbi.nlm.nih.gov/geo/query/acc.cgi?acc=GSM1553879) | Cell Type: Cortex | Sessa A, et al.^3^ |
| NEUROD2 | 74113 | [GSM2253660](http://www.ncbi.nlm.nih.gov/geo/query/acc.cgi?acc=GSM2253660) | Cell Type: Cortex | Guner G, et al.^4^ |
| POLR2A | 86046 | [GSM2803612](http://www.ncbi.nlm.nih.gov/geo/query/acc.cgi?acc=GSM2803612) | Cell Type: Cortex | Stroud H, et al.^5^ |
| TBR1 | 92076 | [GSM3371773](http://www.ncbi.nlm.nih.gov/geo/query/acc.cgi?acc=GSM3371773) | Cell Line: CD-1 | Fazel Darbandi S, et al.^6^ |
| ATAC | - | ENCFF097JFT | e14.5 mouse forebrain | ENCODE Project Consortium.^7^ |

### Supplementary Table 6: Chemicals and reagents

| Reagent | Cat. No. | Brand |
| --- | --- | --- |
| Formaldehyde solution | 28908 | Thermo Fisher Scientific |
| Glycine | 50046 | Sigma-Aldrich |
| DPBS | E607009-0500 | Sangon |
| Nuclease-free Water | AM9930 | Thermo Fisher Scientific |
| Bovine Serum Albumin (BSA) | B2064 | Sigma-Aldrich |
| HEPES pH 7.5 | C0217 | Beyotime |
| NaCl | AM9760G | Thermo Fisher Scientific |
| MgCl_2_ | AM9530G | Thermo Fisher Scientific |
| Spermidine | S0266 | Sigma-Aldrich |
| EDTA-free Protease Inhibitor Cocktail | 04693132001 | Sigma-Aldrich |
| Digitonin | A601152 | Sangon |
| CA-630 | I8896 | Sigma-Aldrich |
| EDTA Solution pH 8.0 | 15575020 | Thermo Fisher Scientific |
| PEG 8000 | R0056 | Beyotime |
| H3K27ac antibody | ab177178 | Abcam |
| H3K27me3 antibody | 39155 | Active Motif |
| H3K4me3 antibody | 39159 | Active Motif |
| Goat Anti-Rabbit IgG H&L | ab6702 | Abcam |
| Goat Anti-Rabbit IgG H&L (Alexa Fluor® 488) | ab150077 | Abcam |
| T4 DNA Ligase | C301-01 | Vazyme |
| T4 DNA polymerase | N101-01 | Vazyme |
| dNTP mix | R0192 | Thermo Fisher Scientific |
| Proteinase K | 19133 | QIAGEN |
| Buffer PKD | 1034963 | QIAGEN |
| NEB buffer 2 | B7002S | New England Biolabs |
| Sodium dodecyl sulfate | 15553027 | Thermo Fisher Scientific |
| Adenosine 5'-Triphosphate (ATP) | P0756S | New England Biolabs |
| Kapa Hotstart HiFi ReadyMix | KK2602 | Roche |
| KAPA SYBR FAST qPCR Master Mix | KK4601 | Roche |
| Equalbit 1 × dsDNA HS Assay Kit | EQ121-01 | Vazyme |
| VAHTS DNA Clean Beads | N411-02 | Vazyme |

### Supplementary Table 7: Summary of sequencing statistics for all samples

| Sample | Sequenced read pairs | Valid barcodes | Spot under tissue | Mean raw read pairs per spot | Confidently mapped read pairs | Fraction of high-quality fragments under tissue | Median high-quality fragments per spot |
| --- | --- | --- | --- | --- | --- | --- | --- |
| E11.5_H3K4me3 | 202708681 | 95.90% | 1564 | 111060 | 89.21% | 87.09% | 23997 |
| E11.5_H3K27me3 | 159287034 | 95.60% | 1591 | 83484 | 78.89% | 85.65% | 15725 |
| E10.5_H3K27ac | 498839562 | 95.50% | 1033 | 386060 | 88.57% | 80.89% | 128804 |
| E11.5_H3K27ac | 565539501 | 96.40% | 1571 | 297639 | 89.10% | 84.90% | 120354 |
| E12.5_H3K27ac | 498463115 | 95.50% | 1474 | 269363 | 88.20% | 82.14% | 77290 |
| E13.5_H3K27ac | 556883708 | 96.20% | 1694 | 278406 | 90.80% | 87.05% | 105424 |
| E14.5_H3K27ac | 523715619 | 95.90% | 1709 | 249414 | 91.77% | 83.80% | 110375 |
| E14.5_Forebrain_H3K27ac | 489990216 | 94.60% | 5390 | 64624 | 95.10% | 76.91% | 15960 |
| E11.5_H3K27ac_slide2 | 69867596 | 95.90% | 1256 | 44849 | 90.67% | 83.14% | 26307 |
| E11.5_H3K4me3_slide2 | 125020414 | 95.50% | 1331 | 81152 | 81.28% | 87.70% | 15686 |
| E11.5_H3K27me3_slide2 | 97808181 | 95.90% | 1162 | 69239 | 77.79% | 82.43% | 12703 |
